# γ-aminobutyrate (GAB) functions as a bioenergetic and signaling gatekeeper to control T cell inflammation

**DOI:** 10.1101/2022.02.17.480926

**Authors:** Siwen Kang, Lingling Liu, Tingting Wang, Matthew Cannon, Penghui Lin, Teresa W.-M. Fan, David A. Scott, Hsin-Jung Joyce Wu, Andrew N. Lane, Ruoning Wang

## Abstract

γ-Aminobutyrate (GAB) is the biochemical form of γ-aminobutyric acid (GABA) at physiological pH and functions as an essential neurotransmitter in the vertebrate’s central nervous system (CNS). Growing evidence suggests that GAB may also mediate intercellular communications to shape various physiological processes, including immune response. Beyond acting as a paracrine signaling molecule, how GAB metabolism is controlled to exert many distinct functions remains elusive. By an integrated analysis of the extracellular metabolome, stable isotope traced metabolic pathway analysis, and metabolic transcriptome, we revealed that GAB is one of the most abundant metabolites produced through glutamine and arginine catabolism in CD4^+^ T help 17 (T_H_17) and induced T regulatory (iT_reg_) cells. GAB functions as a bioenergetic and signaling gatekeeper by reciprocally controlling pro-inflammatory T_H_17 cell and anti-inflammatory iT_reg_ cell differentiation through distinct mechanisms. The expression of 4-aminobutyrate aminotransferase (ABAT) funnels GAB, as an anaplerotic substrate, into the TCA cycle to maximize carbon allocation in promoting T_H_17 cell differentiation. By contrast, the absence of ABAT activities in iT_reg_ cells enables GAB exporting to the extracellular environment and acting as an autocrine signaling metabolite to promote iT_reg_ cell differentiation. Accordingly, genetic or pharmacological ablation of ABAT activity in T cells confers protection against experimental autoimmune encephalomyelitis (EAE) pathogenic progression. Conversely, genetic ablation of GABA(A) receptor in T cells deteriorates EAE pathogenic progression. Collectively, our results suggest that the cell-autonomous control exerted by GAB on CD4^+^ T cell is bimodal and consists of the sequential action of two discrete processes, ABAT–dependent mitochondrial anaplerosis and the receptor-dependent autocrine signaling response, both of which are required for a properly controlled T cell-mediated inflammation.

## Introduction

Mounting a robust and effective adaptive immune response in vertebrates is metabolically costly and requires a proper allocation of essential yet limited energy and carbon resources. Metabolism must be tightly controlled at the cellular level to coordinate a rapid expansion followed by a fine-tuned differentiation process in T cells. Beyond acting as bioenergetic substrates and biosynthetic precursors, metabolites can directly control cellular signaling responses through influencing DNA/RNA/protein modifications, signaling receptors’ activities, and the production of reactive oxygen species ^1–5^. As such, metabolism is fundamental to fine-tuning carbon and nitrogen allocation and optimizing immune response, which is at the center of many diseases. Previous studies have employed systemic approaches to comprehensively characterize the transcriptome, the abundance of intracellular metabolite, and the overall catabolic activities of T cells at the different stages during the T cell life cycle ^6,7^. These studies have generated critical temporal snapshots of the metabolic landscapes, which help establish a conceptual foundation for understanding T cell metabolic reprogramming. However, most of these studies have centered mainly on intracellular metabolites and activities of the central carbon metabolism. The overall metabolic landscape of T cells can also be delineated by monitoring the metabolites consumed from and secreted into the growth medium. The extracellular metabolome represents the ultimate outcome of metabolic input, processing, and output. The extracellular metabolome profiling (also called metabolic footprinting) has been applied as a standard technique to optimize microbial bioprocesses by analyzing substrates consumed from and metabolites secreted into, microorganism’s culture medium ^8,9^. Here, we took a similar approach (**Fig. 1a**) to compare extracellular metabolome profiles of naïve T (T_nai_) cells and different subsets of effector T (T_eff_) cells, including T helper (T_H_0, T_H_1, T_H_17) cells and induced regulator T (iT_reg_) cells.

**Fig. 1.**
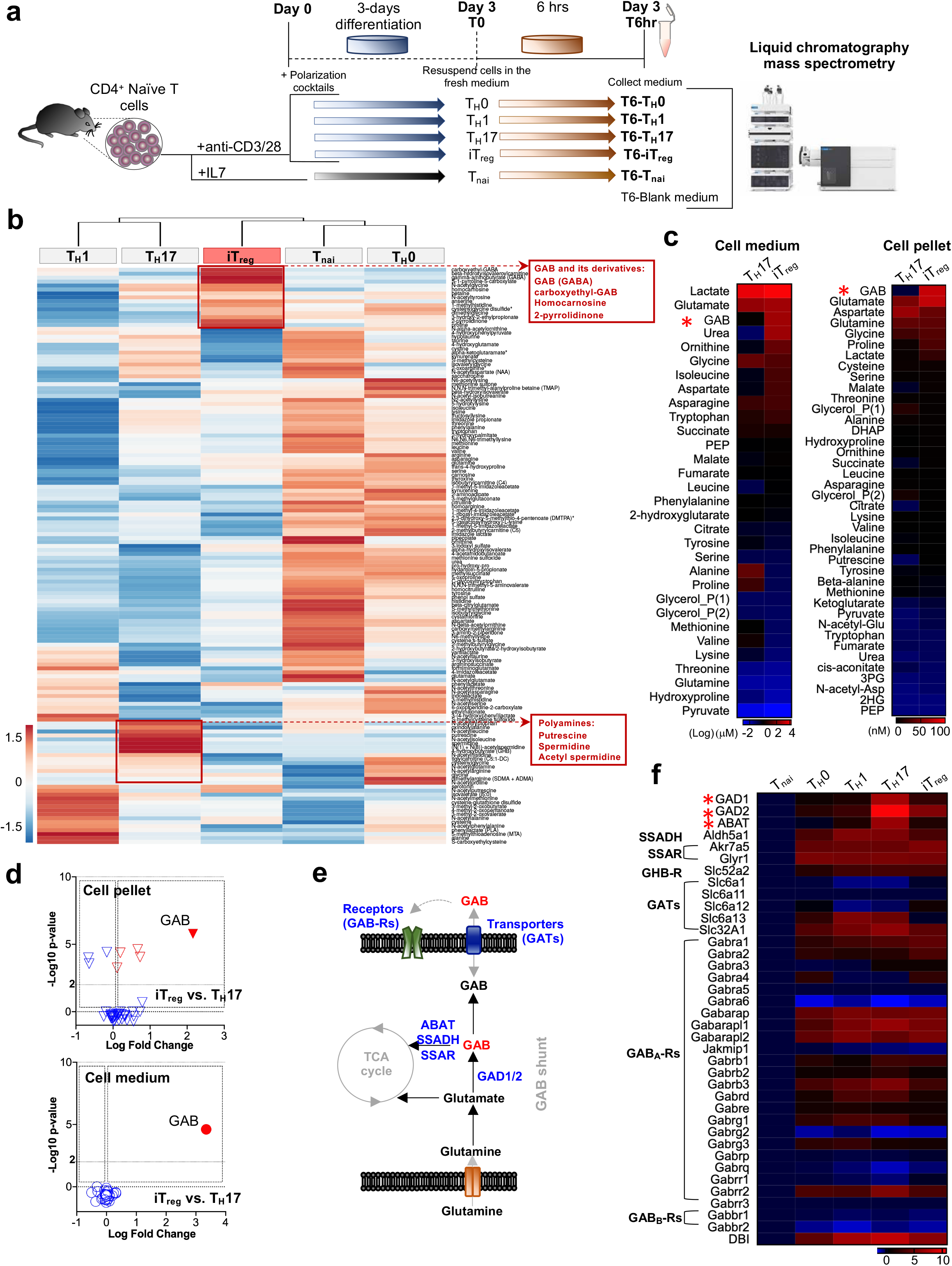
GAB is an abundant metabolite produced in T cells. **a**, Experimental scheme of T cell-extracellular metabolome profiles (LC-MS). **b**, Extracellular metabolites associated with amino acid metabolism in indicated T cell subsets were profiled by LC-MS. The value for each metabolite represents the average of triplicates. The heatmap represents the value of the relative amount (see color scale). The complete metabolomic profile is provided in ***Extended information 1***. **c**, **d**, Indicated metabolites were quantified by GC-MS. The value for each metabolite represents the average of triplicates. The heatmaps (**c**) represent the log2 value (medium) or the absolute value (pellet) of indicated metabolite quantity (see color scale). The complete data is provided in ***Extended information 2***. The volcano plots (**d**) show changes of metabolites in cell pellet (top) and cell medium (bottom). **e,** The schematic pathway of GABA metabolism. **f**, RNA was isolated from indicated T cell subsets (*n*=3) and used for qPCR analyses of indicated metabolic genes. mRNA levels of T_nai_ were set to 1. The heatmap represents the log2 value of the relative mRNA expression level (see color scale). Values and standard deviations are provided in ***Extended information 3***. Statistical analysis was performed by R Programming Language (**b**), unpaired two-tail Student’s *t-*test (**c, d**). Data are pooled from two experiments (**c, d**) or representative of two independent experiments **(f)**. “*n”* indicates the number of biological replicates. *GAD*: glutamate decarboxylase, SSADH: succinic semialdehyde dehydrogenase, SSAR: succinic semialdehyde reductase, GHB-R: γ-hydroxybutyrate receptor, GATs: GABA transporters, VGAT: vesicular GABA transporter, GABA_A_-R: GABA type A receptor, GABA_B_-R: GABA type B receptors, DBI: diazepam binding inhibitor.

## Results

### Extracellular metabolome profiling identifies GAB as an abundant metabolite differentially produced by effector T cells

The control (blank) medium and the spent medium of T cells were profiled on a semi-quantitive untargeted global metabolomics platform based on liquid chromatography-mass spectrometry (LC-MS), with broad coverage of up to 1000-2500 compounds, including amino acids, energy metabolites, nucleotides, and lipids. Using this approach, we have classified metabolites as having changes in production and/or consumption according to whether the fold change compared to control is positive or negative, respectively. The hierarchical clustering analysis, the pairwise comparison, and the principal component analysis revealed that T cell subsets are characterized by distinct extracellular metabolome profiles (**Fig. 1b and S1a-f**). Consistent with the role of the central carbon metabolism in supporting cell growth, the hyper-proliferative T_eff_ groups consume more carbohydrates and produce more lactate than the T_nai_ group (**Fig. S1d**). Also, the T_H_17 group is characterized with the highest production of polyamines among all groups (**Fig. 1b**), in line with the recent finding that revealed a critical role of polyamine in determining T_H_17 differentiation ^10 11,12^. Intriguingly, iT_reg_ cells produce high levels of γ-aminobutyrate (GAB) and its derivatives (**Fig. 1b**). Next, we applied gas chromatograph (GC)-MS-based targeted metabolomics and nuclear magnetic resonance (NMR) to validate and quantify intracellular and extracellular GAB production. We confirmed that iT_reg_ cells produce much higher levels of GAB than T_H_17 cells (**Fig. 1c-d and S2a-b**). Surprisingly, GAB is the most abundant intracellular metabolite and among the top three abundant extracellular metabolites in iT_reg_ cells (**Fig. 1c**). Notably, the intracellular level of GAB is even higher than the level of glutamate (Glu) in iT_reg_ cells, which is one of the most abundant intracellular metabolites in various organisms ^13^. GAB is produced by catabolizing glutamine (Gln) through the GABA shunt and elicits GABAergic response through GABA receptors (GABA-Rs) in neurons. To better understand the molecular nature that determines GAB production and function in T cells, we examined the level of a panel of GABA-related metabolic and receptor genes by qPCR (**Fig. 1e**). Consistent with the previous findings on the GABA-R expression profile in immune cells ^14^. T_eff_ cells express a selected group of GABA-R subunits. However, only iT_reg_ and T_H_17 cells express high levels of glutamate decarboxylase (GAD), the enzyme that catalyzes the decarboxylation of Glu to GAB. Surprisingly, the T_H_17 group exhibits a higher level of GAD than the iT_reg_ group and is the only group that expresses a high level of the GAB catabolizing enzyme, 4-aminobutyrate aminotransferase (ABAT), indicating increased GAB catabolism in T_H_17 cells but not iT_reg_ cells (**Fig. 1f)**. Collectively, these findings suggest that extracellular metabolome profiling is a robust approach to reveal T cell metabolic characters in vitro. Using this approach, we have found that GAB is an abundant metabolite produced by T cells.

### T cells utilize both glutamine and arginine to produce GAB

Given the higher expression of GAD and ABAT in T_H_17 cells than iT_reg_ cells, we reasoned that both iT_reg_ cells and T_H_17 cells could produce GAB. However, the fate of GAB depends on ABAT, i.e., GAB is diverted into the TCA cycle in the presence of ABAT in T_H_17 cells instead of being exported into the extracellular compartment in the absence of ABAT as in iT_reg_ cells. To test this idea, we first validated that ABAT was expressed in T_H_17 cells but not in iT_reg_ cells using Immunoblot (IB) and intracellular staining (**Fig. 2a**). Next, we cultured T_H_17 cells with or without the potent ABAT inhibitor, Vigabatrin (Vig) ^15,16^ for 6 hours and then measured a panel of metabolites. Inhibiting ABAT activities by Vig led to the accumulation of intracellular GAB and GAB release into the medium (**Fig. S2c-d**). Notably, inhibiting ABAT activity renders GAB one of the most abundant metabolites in the medium and pellet (**Fig. S2c-d**). Interestingly, inhibiting ABAT activity reduced Gln consumption without changing Glu levels significantly but increased GAB levels over one-hundred folds (**Fig. 2d**). The reciprocal changes of Gln consumption versus GAB production raise a possibility of a Gln-independent GAB production route in T_H_17 cells. Gln catabolism via the GABA shunt is the canonical GAB biosynthesis pathway ^17^. Alternatively, GAB could be formed from putrescine (Put), a metabolite mainly derived from arginine (Arg) (**Fig. 2c**) ^18,19^. Indeed, the metabolic genes involved in converting arginine into GAB are highly expressed in T_H_17 cells (**Fig. S3a**). To determine to what extent Gln and Arg contribute to GAB biosynthesis, we cultured T_H_17 cells with Vig in the presence or absence of Gln, Arg, or both. Then, we collected spent medium to measure the levels of various metabolites. While removing Gln or Arg reduced GAB production, the removal of both completely blocked GAB production (**Fig. 2e**). Next, we supplied ^13^C_6_-Arg, ^13^C_5_-Gln, ^13^C_6_-Glucose (Glc), or ^13^C_4_-Put as metabolic tracers in culture medium and followed ^13^C incorporation into individual metabolites by GC-MS. The presence of the ^13^C_4_ isotopologue of GAB and the corresponding ^13^C_4_ or ^3^C_5_ isotopologues of upstream metabolites further confirmed that Gln and Arg are carbon donors of GAB (**Fig. 2f-h)**. However, only the ^13^C_2_ isotopologue of GAB was revealed in samples with ^13^C_6_-Glc, suggesting that Glc can support Glu (and GAB) synthesis through the TCA cycle (**Fig. S3b**). Finally, we have shown that Put can be converted to GAB via the diamine oxidase (DAO)-dependent reaction since its inhibitor aminoguanidine (AG) completely blocked the production of ^13^C_4_-GAB from ^13^C_4_-Put (**Fig. S3c**). In addition to a general requirement of both amino acids for protein synthesis, we envisioned that Gln and Arg might also support T_H_17 function and survival through supporting GAB biosynthesis. To test this idea, we cultured T_H_17 cells in the Gln/Arg replete medium or suboptimal medium (with low levels of Gln/Arg) in the absence or presence of high levels of GAB. Supporting our hypothesis, reducing either amino acid led to defects in maintaining viability and IL-17^+^ populations. Notably, the GAB supplement could correct both defects (**Fig. 2i-j)**. We, therefore, conclude that T_H_17 cells can utilize both Gln-derived and Arg-derived carbon to synthesize GAB and support cell viability and function.

**Fig. 2.**
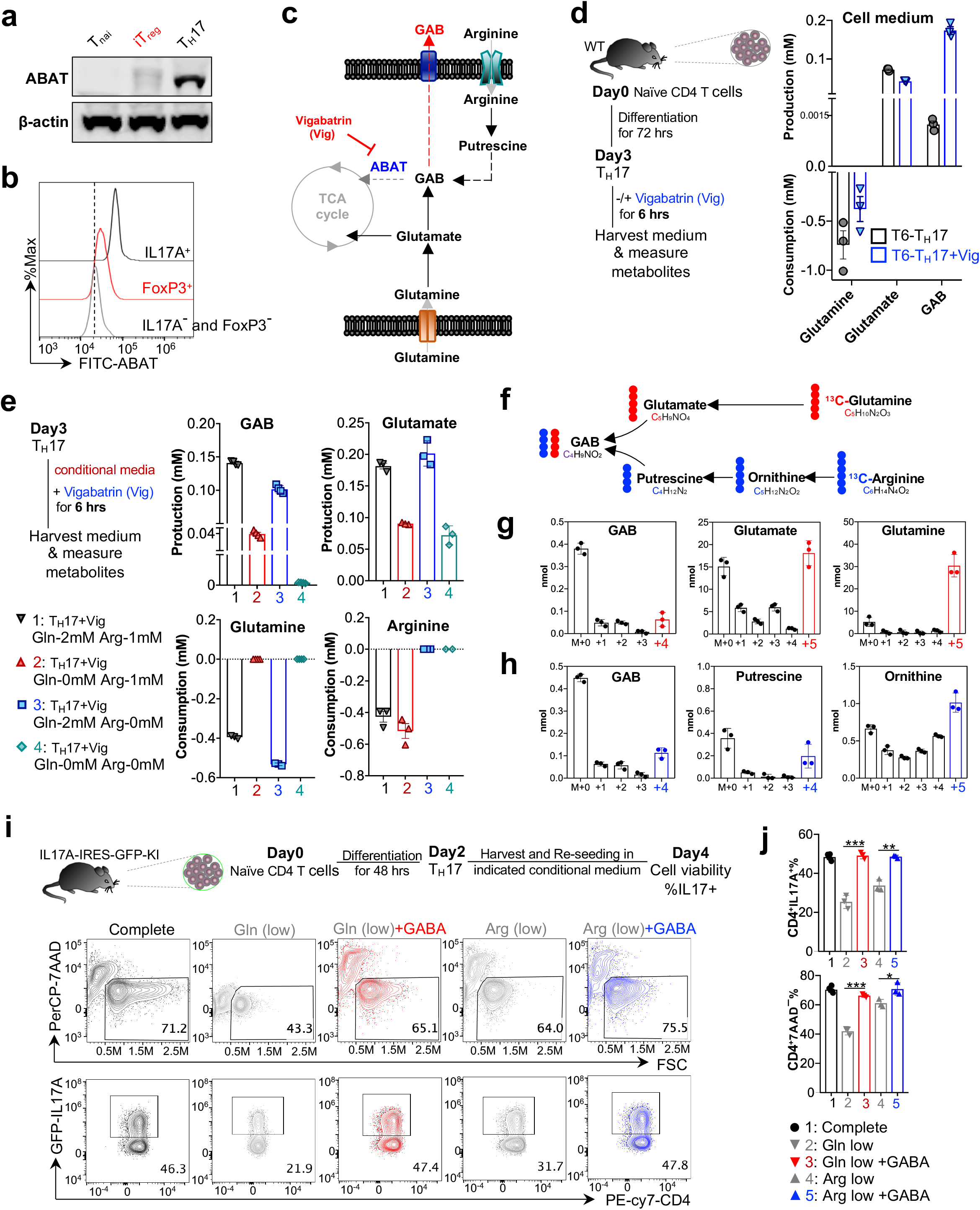
Glutamine and Arginine are the main carbon sources for GAB biosynthesis in effector T cells. **a, b,** ABAT protein levels were determined by **a**, Immunoblot, and **b,** Flow cytometry. **c**, Scheme of GAB metabolic pathways and a pharmacological inhibitor (Vigabatrin) of ABAT. **d**, As illustrated by the experimental scheme (left), indicated metabolites in T_H_17 cells were quantified by GC-MS (right), data are shown as mean a± SEM (*n*=3). **e**, As illustrated by the experimental scheme (left), indicated metabolites in T_H_17 cells were quantified by YSI (Glutamine/ Glutamate) or by bioassay kits (Arginine/ GAB) (*n*=3). **f-h**, Diagram of [^13^C_5_]-glutamine and [^13^C_6_]-arginine conversion to the downstream metabolites (**f**). Indicated metabolites in T_H_17 cells were quantified by GC-MS (*n*=3) (**g, h**). Red or blue dot: ^13^C derived from [^13^C_5_]-glutamine and [^13^C_6_]-arginine. Numbers in the X-axis represent those of ^13^C atoms in given metabolites. **i**, As illustrated by the experimental scheme (top), cytokine expression and cell viability were determined by flow cytometry (bottom). Complete medium: Gln (2 mM), Arg (0.1 mM) and Bicuculline (5 μM), Gln (low): Gln (10 μM), Arg (0.1 mM) and Bicuculline (5 μM), Gln (low plus GABA): Gln (10 μM), Arg (0.1 mM), GABA (1 mM) and Bicuculline (5 μM), Arg (low): Gln (2 mM), Arg (10 μM) and Bicuculline (5 μM), and Arg (low plus GABA): Gln (2 mM), Arg (10 μM), GABA (1 mM) and Bicuculline (5 μM). **j**, Statistical analysis for **i,** data are shown as mean ± SEM (*n*=3). Significance was calculated by Two-way ANOVA (**j**), \**p* < 0.05, \*\**p* < 0.01, \*\*\**p* < 0.001. “*n”* indicates the number of biological replicates. Data are representative of three experiments. Gln: Glutamine, Glu: Glutamate, Arg: Arginine, and MS: mass spectrum.

### ABAT confers on T_H_17 cells a GAB-dependent anaplerosis that maximizes carbon allocation via the TCA cycle

Next, we reasoned that the expression of ABAT may render T_H_17 cells capable of diverting GAB into the TCA cycle in a way to maximize carbon allocation and oxidative phosphorylation (OXPHOS) in mitochondria. To test this idea, we added ^13^C_4_-GABA as a metabolic tracer into the culture medium and followed ^13^C incorporation into intermediate metabolites of the TCA cycle in iT_reg_ cells and T_H_17 cells with or without Vig treatment (**Fig. S4a-b**). In line with the expression of ABAT in T_H_17 cells but not in iT_reg_ cells, T_H_17 cells exhibit much higher levels of ^13^C_4_ isotopologue of succinate and its downstream metabolites in the TCA cycle than iT_reg_ cells (**Fig. S4a)**. Inhibiting ABAT activity by Vig completely abolished ^13^C_4_ isotopologue of succinate and its downstream metabolites in T_H_17 cells, supporting the idea that GAB is diverted to the TCA cycle via ABAT-dependent reaction (**Fig. S4b)**. Next, we sought to determine the temporal change of respiration following a sequential supplement of GABA, Vig, Oligomycin (Oligo), and FCCP into T_H_17 cell culture medium. Indeed, GABA supplement enhanced oxygen consumption via an ABAT-dependent manner, while ATPase inhibitor Oligomycin suppressed and FCCP maximized oxygen consumption as expected (**Fig. S4c)**. We and others have recently shown that Arg-dependent polyamine biosynthesis is required for supporting T cell proliferation and T_H_17 cell differentiation ^10–12^. We reasoned that ABAT expression in T_H_17 cells might allow Arg-derived carbons to be diverted into the TCA cycle through Put and GAB. Supporting this idea, ^13^C_6_-Arg-derived and ^13^C_4_-Put-derived ^13^C were incorporated into the ^13^C_4_ isotopologue of succinate and its downstream metabolites in an ABAT-dependent manner in T_H_17 cells (**Fig. S4d and S5a)**. Finally, we sought to determine if Gln-derived carbons could enter the TCA cycle via ABAT. Gln is a major carbon donor and is known to drive the TCA cycle and OXPHOS via transaminase and glutamate dehydrogenase (GDH) in T_eff_ cells ^20–22^. We found that a sequential supplement of Vig and GDH inhibitor R162 ^23^ suppressed oxygen consumption additively (**Fig. S5c)**. Similarly, the combination of Vig and R162 suppressed ^13^C_5_-Gln-derived ^13^C incorporation into the TCA cycle metabolites more profoundly than single-agent treatment (**Fig. S5b)**. Collectively, we have revealed GAB as a conditional anaplerotic substrate in T cells, and its catabolism via the TCA cycle depends on the expression of ABAT.

### GAB is a metabolic gatekeeper of T cell proliferation and differentiation

To further delineate the role of ABAT in T cells, we generated a T cell-specific *ABAT* knockout strain (*ABAT* cKO) by crossing the *ABAT^fl^* strain with the CD4-Cre strain. qPCR, IB, and intracellular staining analyses validated the deletion of ABAT (**Fig. 3a-b**). ABAT deletion did not result in T cell development defects in the thymus, the spleen, and lymph nodes (**Fig. S6a-f**). In addition, ABAT deletion did not affect cell viability, the expression of cell surface activation markers, the cell cycle progression from G0/G1 to the S phase, RNA/DNA/protein contents, cell size, or viability 24 h after activation in vitro (**Fig. S7a-e**). However, ABAT deletion moderately suppressed overall T cell proliferation after activation in vitro (**Fig. S7e and 3c**). Remarkably, both genetic and pharmacologic ablation of ABAT activity inhibited pro-inflammatory T_H_17 cell differentiation while enhancing anti-inflammatory iT_reg_ cell differentiation *in vitro* (**Fig. 3c**). Supporting these findings, the RNAseq analysis of WT and *ABAT* cKO T cells activated under the T_H_0 condition revealed enriched gene signatures associated with inflammation and T cell differentiation (**Fig. S8a-c**). The expansion and balance between pro-inflammatory CD4^+^ T_eff_ cells and anti-inflammatory CD4^+^ T_reg_ cells determine the pathogenic development of experimental autoimmune encephalomyelitis (EAE), a murine model of multiple sclerosis (MS), which is an inflammatory demyelinating disease of the central nervous system (CNS). Consistent with the expression profile of ABAT in vitro, the IL17^+^ CD4^+^ T group expresses the highest level of ABAT among all the CD4^+^ T subsets infiltration into the central nervous system (CNS) in EAE animals (**Fig. 3d**). Importantly, the genetic deletion of ABAT in T cells or the systemic delivery of Vig conferred significant protection against EAE pathogenic progression, associated with more infiltrated FoxP3^+^ CD4^+^ T cell and less infiltrated inflammatory CD4^+^ T cell, reciprocally (**Fig. 3e-h, and S9a-d**). However, Vig treatment resulted in better protection against EAE and a broader impact on periphery CD4^+^ T cell than T cell-specific deletion of ABAT, indicating that the systemic inhibition of ABAT might affect inflammation through both T cell-intrinsic and T cell-extrinsic mechanisms (**Fig. 3e, 3g, and S9a-d**). We also employed a competitive antigen-specific, TCR-dependent proliferation assay (OT-II) and a competitive homeostatic proliferation assay to assess T cell proliferation and differentiation *in vivo*. Remarkably, the ratio between wild type (WT) and *ABAT* cKO CD4^+^ T cells, CFSE dilution patterns, and the percentage of IL17^+^CD4^+^ T cell in various tissues suggested that the loss of ABAT dampens T cell proliferation and T_H_17 differentiation *in vivo* (**Fig. S10a-e**). Collectively, our results indicate that ABAT status determines the fate of intracellular GAB and, hence, pro-inflammatory T_H_17 and anti-inflammatory iT_reg_ cell differentiation *in vitro* and *in vivo*.

**Fig. 3.**
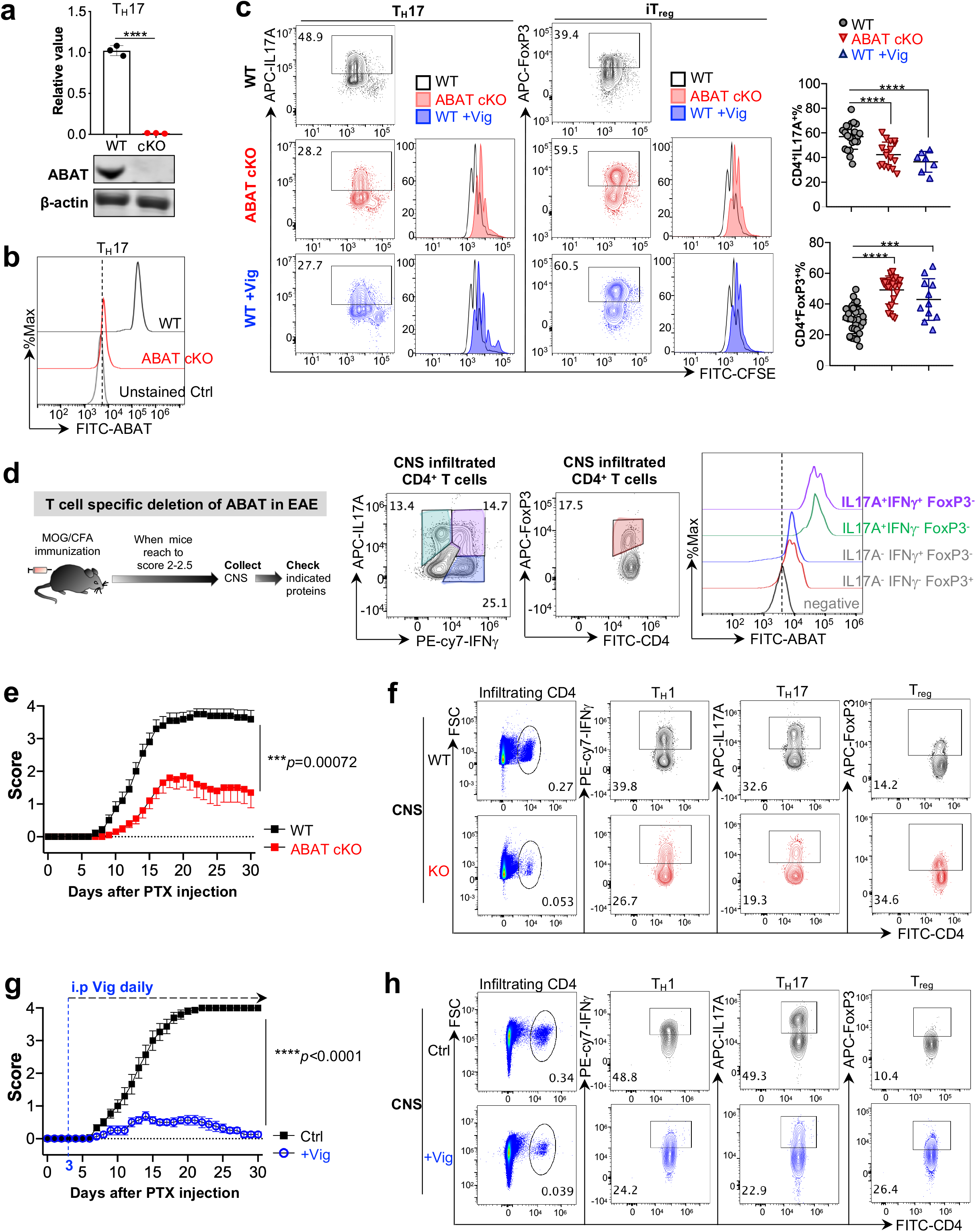
Genetic ablation or pharmacological inhibition of ABAT suppresses T_H_17 but enhances iT_reg_ cell differentiation. **a,b,** ABAT mRNA and protein levels were determined by qPCR and Immunoblot (**a**) or by flow cytometry (**b**). **c**, Expression of indicated cytokines and CFSE dilution in indicated groups were determined by flow cytometry. Statistical analysis of WT (*n*=25), *ABAT* cKO (*n*=15), and WT plus Vig (*n*=7) were shown. **d**, As illustrated by the experimental scheme (left), The expression of indicated proteins in CNS infiltrating CD4^+^ T cells isolated from EAE animals were determined by flow cytometry (*n*=6). **e**, EAE clinical scores in indicated groups were evaluated daily from indicated genotypes (*n*=12/ groups), and **f**, the expression of indicated markers in CNS infiltrating T cells was determined by flow cytometry (*n*=5/ groups). **g**, EAE clinical scores in indicated groups were evaluated daily (*n*=15/ groups), and **h**, The expression of indicated markers in CNS infiltrating T cells were determined by flow cytometry (*n*=3/ groups). Significance was calculated by unpaired Two-tail Student’s *t-*test (**a, e, g**), Two-way ANOVA (**c**), data are shown as mean ± SEM, \*\*\**p* < 0.001, \*\*\*\**p* < 0.0001. “*n”* indicates the number of biological replicates. Data are representative of three experiments (**a**) or pooled from three experiments (**c, e, g**). Vig: vigabatrin, EAE: experimental autoimmune encephalomyelitis, CNS: central nervous system, PTX: paclitaxel, and i.p: intraperitoneal injection.

### GAB can serve as an autocrine signaling metabolite that regulates T cell differentiation through GABA_A_ receptor

In line with earlier studies ^14^, we have found that T_eff_ cells express various subunits of GABA_A_ receptor (GABA_A_-R) (**Fig. 1f**). Also, T cells can produce and secrete a large amount of GAB into the extracellular compartment, which may elicit an autocrine signaling response to regulate T cell differentiation (**Fig. 1c-d, S2a-d, 4a**). Supporting this idea, a low level of GABA supplement could reduce T_H_17 cell but enhance iT_reg_ cell differentiation without affecting T cell activation and proliferation *in vitro* (**Fig. 4b, 4d, S12a, and S12c**). Conversely, GABA_A_-R antagonists with distinct antagonistic mechanisms enhance T_H_17 cell differentiation but reduce iT_reg_ cell differentiation without affecting T cell activation and proliferation *in vitro* (**Fig. 4c, 4e, S12b, and S12d**). The β subunit is a core component of GABA_A_-R, and the β3 subunit (Gabrb3) is highly expressed in all T_eff_ subsets (**Fig. 1f**). We generated a T cell-specific Gabrb3 knockout strain (*Gabrb3* cKO) by crossing the *Gabrb3^fl^* strain with the CD4-Cre strain. Gabrb3 deletion did not result in T cell development defects in the thymus, the spleen, and lymph nodes (**Fig. S11a-d**). In addition, cell viability, the expression of cell surface activation markers, and cell proliferation are comparable in both WT and *Gabrb3* cKO T cells after activation *in vitro* (**Fig. S12e-f**). However, genetic ablation of Gabrb3 promoted pro-inflammatory T_H_17 cell differentiation while reducing anti-inflammatory iT_reg_ cell differentiation *in vitro* (**Fig. 4f**). Importantly, GABA supplement only affects WT but not *Gabrb3* cKO T cell differentiation *in vitro* (**Fig. 4f**). Finally, the T cell-specific *Gabrb3* deletion significantly deteriorates EAE pathogenic progression, associated with increased inflammatory CD4^+^ T cells and decreased FoxP3^+^ CD4^+^ T cells in the CNS and periphery (**Fig. 4g and S13a-b**). Together, these results suggest that GAB is an abundant metabolite produced by T_eff_ cells and exerts both bioenergetic control and autocrine signaling control of T cell differentiation (**Fig. S14**).

**Fig. 4.**
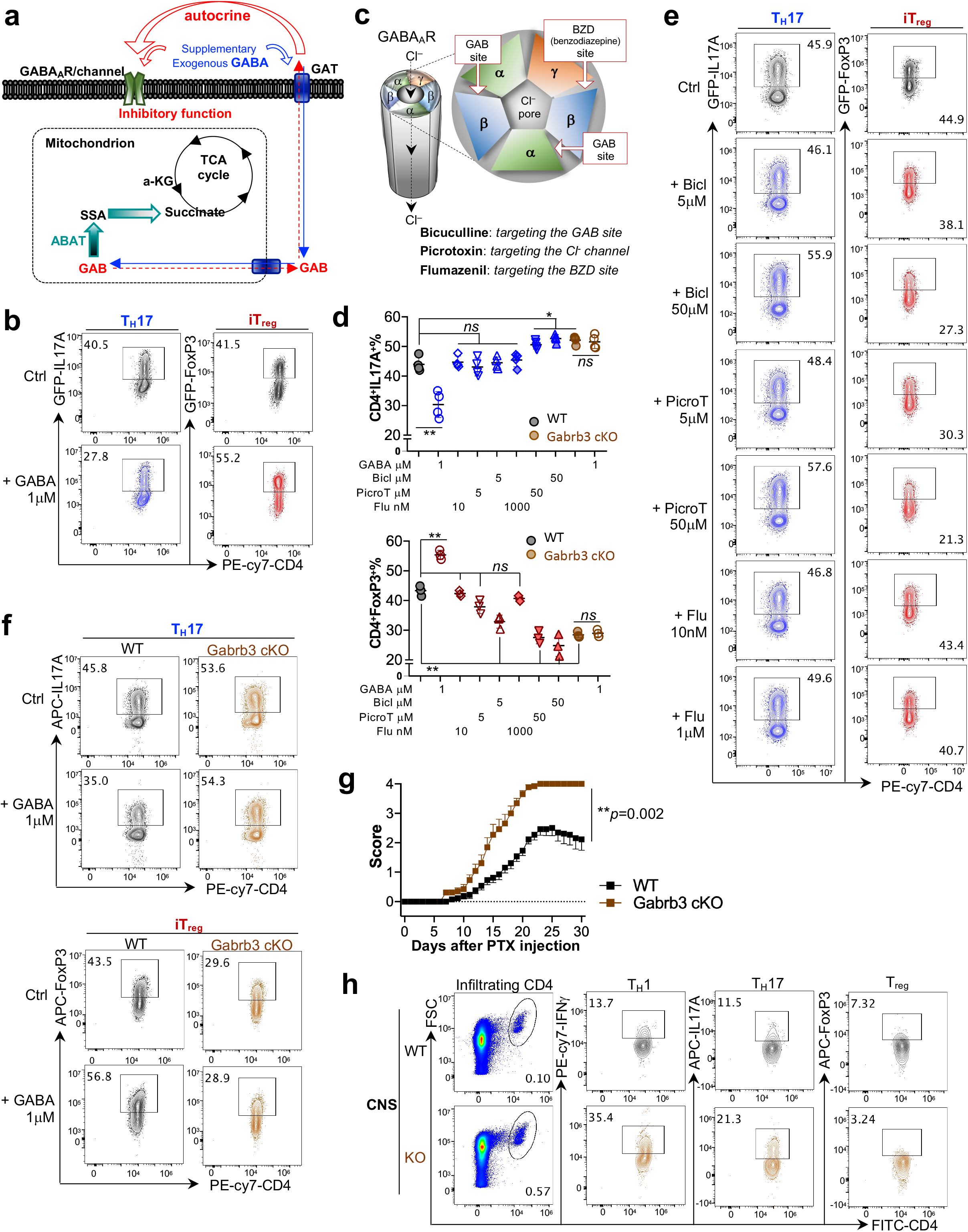
Receptor-mediated GAB autocrine signaling response reciprocally suppresses T_H_17 and enhances iT_reg_ differentiation. **a,** Schematic diagram of GAB metabolism and GABA_A_-R-mediated autocrine signaling response in T cells, and **c**, Schematic diagram of GABA_A_-R antagonists’ binding sites. **b, d, e,** Cytokine expression of indicated groups was determined by flow cytometry. Data are shown as mean ± SEM (*n*=4/ groups). **g**, EAE clinical scores in indicated groups were evaluated daily (*n*=13/ groups), and **h**, The expression of indicated markers in CNS infiltrating T cells were determined by flow cytometry (*n*=3/ groups). Significance was calculated by Two-way ANOVA (**d, combination statistical analysis of b, e, f)**, unpaired Two-tail Student’s *t-*test (**g**), \**p* < 0.05, \*\**p* < 0.01; *ns,* no significant differences. “*n”* indicates the number of biological replicates. Data are representative of two experiments (**b, d, e, f**) or pooled from three experiments (**g**). GABA_A_-R: GABA type A receptor, Cl: chlorine, Bicl: bicuculline, PicroT: picrotoxin, Flu: flumazenil, EAE: experimental autoimmune encephalomyelitis, CNS: central nervous system, and PTX: paclitaxel

## Discussion

The vertebrate immune and nervous systems are intimately connected with each other developmentally, anatomically, and physiologically. Interaction between the two systems coordinates their sensory functions to ensure organismal homeostasis and survival ^24,25 26 27^. Immune cells and neurons can communicate with each other through a group of shared ligand molecules and receptors, including the neurotransmitter GABA and its receptors ^14,28^. Beyond mediating intersystem communication between the immune and nervous system, growing evidence suggests that GABA can also act as a paracrine signaling molecule mediating intrasystem communication to regulate immune response ^29^. One recent study has found that B cells can produce GABA and suppress anti-tumor immunity through paracrine modulation of intratumoral macrophages and CD8^+^ T cells ^30^. Also, GABA in macrophages has been implicated as an intracellular metabolite with a pro-inflammatory function ^14^. Here we show that GAB (the biochemical form of GABA at physiological pH) is one of the most abundant metabolites in T cells and promotes inflammation through modulating T cell proliferation and differentiation. Depending on the status of its catabolizing enzyme ABAT, GAB can act as a conditional anaplerotic substrate to promote T_H_17 cell differentiation or an autocrine signaling metabolite to enhance iT_reg_ cell differentiation. In addition to its role in mediating intercellular communications, GAB also serves as a metabolic and signaling gatekeeper to regulate inflammation through a T cell-autonomous manner.

T_eff_ cells consume glutamine and arginine at high rates ^31,32^. Beyond a general requirement for protein synthesis, glutamine and arginine support T cell proliferation and function through their catabolic products. Glutamine is a primary carbon source to sustain the TCA cycle, which generates energy through oxidative phosphorylation and allocates carbon to produce biosynthetic precursors to support T cell growth ^21,32,33^. Similarly, arginine catabolism is coupled with the urea cycle to produce bioactive metabolites such as polyamines to support T cell proliferation and differentiation ^10–12^. Our results show that both arginine catabolism (via putrescine) and glutamine catabolism (via glutamate) are coupled with GAB biosynthesis in T_H_17 cells, implicating GAB as a crucial metabolic node and a branch point in amino acids catabolism. GAB can be consumed through the TCA cycle to enhance bioenergetic and biosynthetic capacities or be secreted as an autocrine signaling metabolite depending on the status of ABAT. We have further revealed that glutamine can replenish the TCA cycle intermediate metabolites through either glutamate or GAB anaplerosis. Glutamate increases the α-KG level while GAB increases the succinate level. Therefore, it is conceivable that the carbon input from glutamate or GAB may change the intracellular α-KG /succinate ratio reciprocally. Hence, the GABA shunt in T cells may impact the hypoxia signaling response and/or DNA/histone methylation pattern through modulating the enzymatic activities of the α-KG-dependent dioxygenase family ^33–35^. Notably, glutamate and putrescine are highly abundant intracellular metabolites that can be secreted to the extracellular environment by T_H_17 cells (**Fig 1a**) ^10,36^. The GAB catabolizing enzyme, ABAT, may provide a sensitive and precise regulation of the three interconnected and highly abundant metabolites: GAB, glutamate, and putrescine, permitting rapid metabolic and signaling responses to control inflammation.

T cells’ high and dynamic metabolic demand during inflammatory and autoimmune responses requires fine-tuned central carbon and ancillary metabolic pathways regulation. Hence, metabolic pathways have been therapeutically exploited to target inflammatory and autoimmune diseases ^37,38^. Disruption of the central carbon catabolism can affect many cellular processes and cell types. However, targeting ancillary metabolic pathways that are only engaged in a small group of specialized immune cells under physio-pathological conditions may result in less toxicity but maximal clinical benefits ^6^. Gene and protein expression profiling studies have suggested that human autoimmune diseases, including multiple sclerosis (MS), type 1 diabetes, and rheumatoid arthritis, are associated with the dysregulation of GABA-related metabolic and signaling genes ^39–42^. Interestingly, the cortical GAB levels are lower in patients with relapsing-remitting multiple sclerosis (RRMS) than healthy controls ^43,44^. In addition, one recent study based on genome-scale metabolic modeling and in silico simulations for drug response indicates that GAB metabolism and signaling pathway are not only involved in the disease process but also are potential drug targets in human autoimmune diseases ^45^. Consistent with clinical profiling and in silico studies, pharmacological modulation of GAB metabolism and receptor-mediated signaling response could ameliorate pathological phenotypes in several preclinical models of autoimmune diseases ^46–50^. Our results further elucidate a previously unrecognized aspect of the T cell-intrinsic effects conferred by GAB catabolism and receptor-mediated signaling. Collectively, GAB modulating strategies via blockade of GAB catabolism, activation of receptor-mediated response, or both may present a promising and novel therapy for treating inflammatory and autoimmune diseases.

## Data availability statement

The RNA-seq datasets generated for this study can be found in the GEO accession GSE190818.

## Materials and Methods

### Mice

C57BL/6 (WT), Flippase (B6.129S4*Gt(ROSA)26Sor^tm1(FLP1)Dym^*/RainJ), OT-II (B6.Cg-Tg(TcraTcrb)425Cbn/J), CD45.1^+^ (B6.SJL-*Ptprc^a^Pepc^b^*/BoyJ), *Rag1*^-/-^ (B6.129S7-*Rag1^tm1Mom^*/J), IL17A-IRES-GFP-KI (C57BL/6-*Il17a^tm1Bcgen^*/J), FoxP3^GFP+^ (C57BL/6-Tg(Foxp3-GFP)90Pkraj/J), and Gabrb3*^fl^* (B6;129-*Gabrb3^tm2.1Geh^*/J) mice were obtained from the Jackson Laboratory (JAX, Bar Harbor, ME). Mice with one targeted allele of *ABAT* on the C57BL/6 background (*ABAT*^tm1a(EUCOMM)Hmgu^) were generated by The European Conditional Mouse Mutagenesis Program (EUCOMM) ^51^. The mice were first crossed with a transgenic Flippase strain (B6.129S4*Gt(ROSA)26Sor^tm1(FLP1)Dym^*/RainJ) to remove the LacZ-reporter allele and then crossed with the CD4-Cre strain to generate T cell-specific *ABAT* knockout strain (*ABAT* cKO). OT-II mice were crossed with CD4Cre *ABAT* cKO mice to generate the OT- II CD4Cre *ABAT* cKO mice. OT-II mice were crossed with *Thy1.1*^+^ mice (B6.PL-*Thy1*^a^/CyJ) to generate the OT-II *Thy1.1* mice. Gabrb3*^fl^* mice were crossed with the CD4-Cre strain to generate T cell-specific *Gabrb3* knockout strain (*Gabrb3* cKO). Gender and age-matched mice (6-12 weeks old) were used in the experiments. All mice were bred and kept in specific pathogen-free conditions at the Animal Center of Abigail Wexner Research Institute at Nationwide Children Hospital. Animal protocols were approved by the Institutional Animal Care and Use Committee of the Abigail Wexner Research Institute at Nationwide Children’s Hospital (IACUC; protocol number AR13-00055).

### Murine T cell Isolation and Culture

Naïve CD4^+^ T cells were enriched from mouse spleen and lymph nodes by negative selection using MojoSort Mouse CD4 Naive T Cell Isolation Kit (MojoSort, Biolegend) following the manufacturer’s instructions. For the activation assay, freshly isolated CD4^+^ T cells were either maintained in culture medium with 5 ng/mL IL-7 for resting-state or were activated with 5 ng/mL IL-2 and plate-bound anti-mCD3 and anti-mCD28. The culture plates were pre-coated with 2 μg/mL anti-mCD3 and 2 μg/mL anti-mCD28 antibodies overnight at 4°C. Unless indicated separately, the cells were seeded in the RPMI-1640 medium supplemented with 10% FBS, or (v/v) heat-inactivated dialyzed fetal bovine serum (DFBS), 2 mM L-glutamine, 1% sodium pyruvate (Sigma-Aldrich), 100 units/mL penicillin, 100 μg/mL streptomycin, and 0.05 mM 2-mercaptoethanol (Sigma-Aldrich) at 37°C/5% CO_2_.

For CD4 T cell differentiation, 48 wells culture plates were pre-coated with 2 μg/mL (iT_reg_ differentiation), 5 μg/mL (T_H_1 differentiation) or 10 μg/mL (T_H_17 differentiation) anti-mCD3 and anti-mCD28 antibodies over nights at 4°C. Freshly isolated naïve CD4^+^ T cells (0.5 ×10^6^/mL) were activated with plate-bound antibodies and with mIL-2 (3 ng/mL) and hTGF-β1 (10 ng/mL) for iT_reg_ differentiation), or with mIL-2 (10 ng/mL) and mIL-12 (20 ng/mL) for T_H_1 differentiation, or with mIL-6 (50 ng/mL), hTGF-β1 (20 ng/mL), anti–mIL-2 (8 μg/mL), anti– mIL-4 (8 μg/mL), and anti–mIFN-γ (8 μg/mL) for T_H_17 differentiation. In some experiments, Vigabatrin (Vig, 1 mM), the γ-Aminobutyric acid (GABA, 0.1 μM~1 mM), GABA_A_-R antagonists including Bicuculline (Bicl, 5 μM, 50 μM), Picrotoxin (PicroT, 5 μM, 50 μM), and Flumazenil (10 nM, 1 μM), R162 (20 μM), Oligomycin (1.5 μM), FCCP (1 μM), or aminoguanidine (AG, 0.2 mM) was added to cell culture medium. Additional information of cytokines, antibodies, and chemicals were list in **Table 1**.

### Flow Cytometry

For analyzing surface markers, cells were stained in PBS containing 2% (w/v) BSA and the appropriate antibodies from Biolegend. For analyzing intracellular cytokine IFN-γ and IL-17A, T cells were stimulated for 4 hrs with eBioscience™ Cell Stimulation Cocktail (eBioscience) before being stained with cell-surface antibodies. Cells were then fixed and permeabilized using FoxP3 Fixation/Permeabilization solution according to the manufacturer’s instructions (eBioscience). Cell proliferation was assessed by CFSE staining per the manufacturer’s instructions (Invitrogen). Cell viability was evaluated by 7AAD staining per the manufacturer’s instructions (Biolegend). For analyzing DNA/RNA content, cells were collected and stained with surface markers before being fixed with 4% paraformaldehyde for 30 min at 4°C, followed by a step of permeabilization with FoxP3 permeabilization solution (eBioscience). Cells were stained with 7AAD for 5 min and then stained with pyronin-Y (4 μg/ml, PE) for 30 min before being analyzed by flow cytometer with PerCP channel for 7AAD (DNA) and PE channel for pyronin-Y (RNA). Protein synthesis assay kit (Item No.601100, Cayman) was used for analyzing protein content. Briefly, cells were incubated with O-propargyl-puromycin (OPP) for 1 hr, then were fixed and stained with 5 FAM-Azide staining solutions before being analyzed by flow cytometer with FITC channel. For analyzing cell cycle profile, cells were incubated with 10 μg/mL BrdU for 1 hr, followed by cell surface staining, fixation, and permeabilization according to Phase-Flow Alexa Fluro 647 BrdU Kit (Biolegend). Flow cytometry data were acquired on Novocyte (ACEA Biosciences) and were analyzed with FlowJo software (TreeStar). Additional information on flow cytometry antibodies was listed in **Table 2.**

### Western Blot Analysis

For protein extraction, cells were lysed and sonicated at 4°C in a lysis buffer (50 mM Tris-HCl, pH 7.4, 150 mM NaCl, 0.5% SDS, 5 mM sodium pyrophosphate, protease, and phosphatase inhibitor tablet), then centrifuged at 13,000 ×*g* for 15 min for recovering. The samples were boiled in the mixture of LDS Sample Buffer (NuPAGE) and Reducing solution (Thermo Fisher Scientific) for 5 min, after transferring to PVDF membranes by using the iBlot Gel Transfer Device (Thermo Fisher Scientific), then incubated with primary anti-ABAT (clone B-12, Santa Cruz Biotechnology) followed by incubating with the secondary antibodies conjugated with horseradish peroxidase. Immunoblots were developed on films using the enhanced chemiluminescence technique.

### RNA Extraction, qPCR, and RNAseq

Total RNA was isolated by using the RNeasy Mini Kit (Qiagen). The cDNA synthesis was processed using Random hexamers and M-MLV Reverse Transcriptase (Invitrogen). BIO-RAD CFX284^TM^ Real-Time PCR Detection System was used for SYBR green-based quantitative PCR. The relative gene expression was determined by the comparative *CT* method, also referred to as the 2^−ΔΔ*CT*^ method. The data were presented as the fold change in gene expression normalized to an internal reference gene (beta2-microglobulin) and relative to the control (the first sample in the group). Fold change = 2^−ΔΔ*C*^T = [(*CT* _gene of interest_-*CT* _internal reference_)] sample A - [(*CT* _gene of interest_-*CT* _internal reference_)] sample B. Samples for each experimental condition were run in triplicated PCR reactions. Primer sequences were obtained from Primer Bank to detect target genes (**Table 3**).

For RNA sequencing analysis, total RNA was extracted using RNeasy Mini Kit (Qiagen) and treated with DNase I according to the manufacturer’s instructions. After assessing the quality of total RNA using an Agilent 2100 Bioanalyzer and RNA Nanochip (Agilent Technologies), 150 ng total RNA was treated to deplete the levels of ribosomal RNA (rRNA) using target-specific oligos combined with rRNA removal beads. Following rRNA removal, mRNA was fragmented and converted into double-stranded cDNA. Adaptor-ligated cDNA was amplified by limit cycle PCR. After library quality was determined via Agilent 4200 TapeStation and quantified by KAPA qPCR, approximately 60 million paired-end 150 bp sequence reads were generated on the Illumina HiSeq 4000 platform. Quality control and adapter trimming were accomplished using the FastQC (version 0.11.3) and Trim Galore (version 0.4.0) software packages. Trimmed reads were mapped to the Genome Reference Consortium GRCm38 (mm10) murine genome assembly using TopHat2 (version 2.1.0), and feature counts were generated using HTSeq (version 0.6.1). Statistical analysis for differential expression was performed using the DESeq2 package (version 1.16.1) in R, with the default Benjamini-Hochberg *p-value* adjustment method. The Ingenuity Pathway Analysis (IPA) software (QIAGEN), the Gene Set Enrichment Analysis (GSEA) software (UC San Diego, BROAD Ins.), and the R Programming Language software were used to analyze gene signature and pathway enrichment.

### Adoptive Cell Transfer assays

For homeostatic proliferation in lymphopenic *Rag^-/-^* mice, naïve CD4^+^ T cells isolated from donor mice using naïve CD4^+^ mouse T cell isolation kit (Biolegend) were labeled with CFSE. Approximately 1 ×10^7^ cells (mixed with WT and KO cells at 1:1 ration) in 150 μL PBS were transferred via caudal venous injection into 6-8-week-old gender-matched host mice. Mice were sacrificed between 4-7 days after cell transfer. Lymph nodes and spleen were collected and processed to assess cell ratio and proliferation by flow cytometry analysis.

For antigen-driven proliferation using OTII mice: naïve CD4^+^ T cells isolated from OTII/CD45.2 TCR transgenic donor mice using naïve CD4^+^ mouse T cell isolation kit (Biolegend) were labeled with CFSE. Approximately 1 ×10^7^ cells (mixed with WT and KO cells at 1:1 ration) in 150 μL PBS were transferred via caudal venous injection into 6-8-week-old gender-matched CD45.1 host mice. Host mice were immunized subcutaneously in the hock area (50 μL each site) in both legs with 1 mg/mL OVA^323-339^ peptide (InvivoGen) emulsified with CFA (InvivoGen). Mice were sacrificed 8 days after immunization. Lymph nodes were collected and processed to assess cell ratio, proliferation, and protein expression by flow cytometry analysis.

### Experimental Autoimmune Encephalomyelitis (EAE)

Mice were immunized subcutaneously with 100 μg of myelin oligodendrocyte glycoprotein (MOG)_35–55_ peptide emulsified in complete freund adjuvant (CFA), which was made from IFA(Difco) plus mycobacterium tuberculosis (Difco). Mice were i.p. injected with 200 ng of pertussis toxin (PTX, List Biological Laboratories) on the day of immunization and 2 days later. In experiments as shown in **Fig 3g, 3h, and S9c-d**, mice were i.p injected with 250 mg/kg in 100 μL PBS daily since day 3 after immunization throughout the experiment. In experiments as shown in **Fig 4g, 4h, and S13**, animals were injected with PTX only once on the day of immunization for a sub-optimal EAE induction. All mice were observed daily for clinical signs and scored as described previously ^10^. In some experiments, mice were euthanized when control mice reached the onset of symptoms. The CNS (brain and spinal cord), spleen, and peripheral lymph nodes were collected and mashed to make the single-cell solution. The cell suspension was centrifuged on a 30%/70% Percoll gradient at 500 *g* for 30 min to isolate mononuclear cells from the CNS, followed by cell surface and intracellular staining and flow cytometric analysis described above.

### Stable Isotope Labeling Experiments

#### ^13^C_5_-Glutamine, ^13^C_6_-Arginine, and ^13^C_6_-Glucose labeling of T_H_17 cells (see Fig 2f-h and S3b)

Naïve CD4 T cells isolated from WT mice were polarized for 72 hrs under T_H_17 culture condition before being collected and re-seeded at the density of 2 ×10^6^ cells /mL in conditional medium (RPMI-1640) containing 4 mM ^13^C_5_-Glutamine, 1 mM ^13^C_6_-Arginine, or 10 mM ^13^C_6_-Glucose. After 12 hrs of culture, around ~1 ×10^7^ cells for each sample were collected and washed 3 times with PBS before being snap-frozen.

#### ^13^C_6_-Arginine labeling of T_H_17 cells (see S4d)

T_H_17 cells (as described above) were pretreated with vehicle or Vig (1 mM) for 1hr before being collected and re-seeded at the density of 2 ×10^6^ cells /mL in conditional medium (RPMI-1640) containing 4 mM 1 mM ^13^C_6_-Arginine with vehicle or Vig (1 mM). After 6 hrs of culture, around ~1 ×10^7^ cells for each sample were collected and washed 3 times with PBS before being snap-frozen.

#### ^13^C_4_-Putruscine labeling of T_H_17 cells (see S3c and S5a)

T_H_17 cells (as described above) were pretreated with vehicle, Vig (1 mM) or AG (0.2 mM) for 1hr, and then were collected and re-seeded at the density of 2 ×10^6^ cells /mL in conditional medium (RPMI-1640) containing 0.1 mM ^13^C_4_-Putruscine and 10 μM Arginine and with vehicle, Vig (1 mM) or AG (0.2 mM) treatment. After 6 hrs of culture, around ~1 ×10^7^ cells for each sample were collected and washed 3 times with PBS before being snap-frozen.

#### ^13^C_4_-GABA labeling of T_H_17 and iT_reg_ cells (see S4a)

Naïve CD4 T cells isolated from WT mice were polarized for 72 hrs under T_H_17 or iT_reg_ culture condition before being collected and re-seeded at the density of 2 ×10^6^ cells /mL in conditional medium (RPMI-1640) containing 0.5 mM ^13^C_4_-GABA, 0.1 mM glutamine, and GABA_A_-R antagonist Bicuculline (5 μM). After 12 hrs of culture, around ~1 ×10^7^ cells for each sample were collected and washed 3 times with PBS before being snap-frozen.

#### ^13^C_4_-GABA labeling of T_H_17 with Vig (see S4b)

T_H_17 cells (as described above) were pretreated with vehicle or Vig (1 mM) for 1hr before being collected and re-seeded at the density of 2 ×10^6^ cells /mL in the conditional medium containing 0.5 mM ^13^C_4_-GABA, 0.1 mM glutamine, and GABA_A_-R antagonist Bicuculline (5 μM) and with vehicle or Vig (1 mM) treatment. After 6 hrs of culture, around ~1 ×10^7^ cells for each sample were collected and washed 3 times with PBS before being snap-frozen.

#### ^13^C_5_-Glutamine labeling of T_H_17 cells with multiple inhibitors (see S5b)

T_H_17 cells (as described above) were pretreated with vehicle, Vig (1 mM) or R162 (20 μM) for 1hr, and then were collected and re-seeded at the density of 2 ×10^6^ cells /mL in conditional medium (RPMI-1640) containing 4 mM ^13^C_5_-Glutamine and with the vehicle, Vig (1 mM), R162 (20 μM) or the combination of Vig and R162 treatment. After 6 hrs of culture, around ~1 ×10^7^ cells for each sample were collected and washed 3 times with PBS before being snap-frozen.

Additional information on stable isotope items was listed in **Table 4**.

### Gas Chromatography-Mass Spectrometry (GC-MS) Sample Preparation and Analysis

GC-MS was performed as previously described ^52^, cell pellets were resuspended in 0.45 mL −20°C methanol/ water (1:1 v/v) containing 20 μM L-norvaline as internal standard. Further extraction was performed by adding 0.225 mL chloroform, vortexing, and centrifugation at 15,000 ×*g* for 5 min at 4°C. The upper aqueous phase was evaporated under vacuum using a Speedvac centrifugal evaporator. Separate tubes containing varying amounts of standards were evaporated. Dried samples and standards were dissolved in 30 μL 20 mg/mL isobutylhydroxylamine hydrochloride (TCI #I0387) in pyridine and incubated for 20 min at 80°C. An equal volume of N-tertbutyldimethylsilyl-N-methyltrifluoroacetamide (MTBSTFA) (Soltec Ventures) was added and incubated for 60 min at 80°C. After derivatization, samples and standards were analyzed by GC-MS using a Rxi-5ms column (15 m × 0.25 i.d. × 0.25 μM, Restek) installed in a Shimadzu QP-2010 Plus gas chromatograph-mass spectrometer (GC-MS). The GC-MS was programmed with an injection temperature of 250°C, 1.0 μL injection volume, and a split ratio of 1/10. The GC oven temperature was initially 130°C for 4 min, rising to 250°C at 6°C/min, and to 280°C at 60°C/min with a final hold at this temperature for 2 min. GC flow rate, with helium as the carrier gas, was 50 cm/s. The GC-MS interface temperature was 300°C, and (electron impact) ion source temperature was 200°C, with 70 eV ionization voltage. Fractional labeling from ^13^C substrates and mass isotopomer distributions were calculated as described ^52^. Data from standards were used to construct standard curves in MetaQuant ^53^, from which metabolite amounts in samples were calculated. Metabolite amounts were corrected for recovery of the internal standard and for ^13^C labeling to yield total (labeled and unlabeled) quantities in nmol per sample and then adjusted per cell number.

### Liquid Chromatography-Mass Spectrometry (LC-MS) Sample Preparation and Analysis

Naïve CD4^+^ T cells were polarized under T_H_0, T_H_1, T_H_17, and iT_reg_ culture conditions or cultured with IL-7 (T_nai_ condition) for 72 hrs. Then, cells were harvested, washed by PBS, and then re-seeded at 5 ×10^6^ cells /mL density in the fresh medium. After 6 hrs culture, the cell medium was collected and snap-frozen. Sample preparation and analysis were carried out as described previously at Metabolon, Inc ^54^. In brief, sample preparation involved protein precipitation and removal with methanol, shaking, and centrifugation. The resulting extracts were profiled on an accurate mass global metabolomics platform consisting of multiple arms differing by chromatography methods and mass spectrometry ionization modes to achieve broad coverage of compounds differing by physiochemical properties such as mass, charge, chromatographic separation, and ionization behavior. Metabolites were identified by automated comparison of the ion features in the experimental samples to a reference library of chemical standard entries that included retention time, molecular weight (*m/z*), preferred adducts, and in-source fragments as well as associated MS spectra, and were curated by visual inspection for quality control using software developed at Metabolon.

### NMR analysis of medium

Naïve CD4^+^ T cells isolated from WT mice were polarized for 72 hrs under T_H_17 or iT_reg_ culture conditions. Cells were harvested, washed by PBS, and then re-seeded at a 2 ×10^6^ cells /mL density in conditional medium (RPMI-1640) containing 4 mM ^13^C_5_-Glutamine. Medium samples were taken at 0 hr, and after 6 hrs incubation, were extracted and lyophilized. The dried samples were reconstituted in deuterated 50 mM phosphate buffer pH 8 containing 8.81 nmole DSS-d_6_. NMR spectra were recorded on a Bruker Avance III spectrometer at 16.45 T at 15 °C in a 1.7 mm HCN inverse triple resonance cryoprobe. Presat spectra were recorded with an acquisition time of 2 s with weak irradiation at the HOD frequency during the relaxation delay of 4 s (pulse program: ZGPR). ^1^H{^13^C} HSQC spectra were recorded with an acquisition time of 0.25 s and a relaxation delay of 1.75 using adiabatic decoupling of ^13^C (pulse program: hsqcetgpprsisp2.2). The raw data were apodized with a 1 Hz exponential line broadening and linear predicted once (Presat) or a 4-Hz line broadening exponential (HSQC), phased, and baseline corrected using third-order Bernstein polynomials and referenced to the internal DSS at 0 ppm. Peaks were deconvoluted by line fitting using MNOVA and normalized to the DSS resonance at 0 ppm. The HCCH-TOCSY 2D spectrum was acquired by recording the first plane of the 3D experiment in the carbon dimension (pulse program: hcchdigp3d). The acquisition times in the direct and indirect dimensions (f_2_ and f_1_) are 0.25 s and 0.03 s, respectively. The C-C mixing time was set to 10.9 ms using a DIPSI-3 spin lock scheme with B_1_ field strength of 40 kHz. The data were processed with 1 Hz exponential apodization in f_2_ and 5 Hz exponential with cosine squared function in the f_1_ dimension, and further linear predicted and zero-filled to 8 k and 2k data points, respectively. The spectrum was phased and baseplane corrected before referencing to glutamine peaks based on standard spectrum. Spectral assignments were made by reference to authentic standards of GAB recorded under the same conditions.

### Metabolite quantification

In some experiments, T_H_17 cells were suspended at the density of 5 ×10^6^ cells /mL with vehicle or Vig (1 mM). After 6 hrs culture, blank media (without cells) and spent media were collected. The levels of glutamine and glutamate were measured by the bioanalyzer (YSI 2900). According to the manufacturer’s instructions, arginine and GAB were determined by L-Arginine Assay Kit (BioVision) and GABA Research ELISA Kit (LDN). Each metabolite consumption or production was determined by calculating the difference between blank and spent media.

### Oxygen consumption rate (OCR)

According to the manufacturer’s instruction, OCR was determined by using seahorse XFe96 Analyzer (Agilent Technologies). Briefly, approximate ~1 ×10^5^ T_H_17 cells were suspended in 50 μL assay medium (Seahorse XF RPMI Assay Medium, pH 7.4, Agilent Technologies) containing 10 mM glucose, 2 mM glutamine, and 1 mM pyruvate were seeded in a Poly-D-Lysine (50 μg/ mL, Millipore) pre-coated XF96 Cell Culture Microplates (Seahorse, Agilent Technologies). Cells were centrifuged at 200 ×*g* for 2 min in a zero-braking setting to immobilize cells before being supplied with an additional 130 μL assay medium and kept in a non-CO_2_ incubator for 30 min. Data analysis was performed using the Seahorse Wave Software (Seahorse, Agilent Technologies). In some experiments, the GABA_A_-R antagonist Bicuculline (5 μM) was added along with GABA to prevent the activation of GABA_A_-R. Various compounds were injected into each well sequentially to achieve the following final concentrations: 0.5 mM (GABA), 1 mM (Vig), 20 μM (R162), 1.5 μM (Oligomycin), and 1 μM (FCCP).

### Statistical Analysis

Statistical analysis was conducted using the GraphPad Prism software (GraphPad Software, Inc.). *P* values were calculated with Two-way ANOVA for the EAE experiments. Unpaired two-tail Student’s *t*-test, multiple comparisons of One/ Two-way ANOVA were used to assess differences in other experiments. R Programming Language software was used for Metabolon and RNA-seq data analysis. *P* values smaller than 0.05 were considered significant, with *p* < 0.05, *p* < 0.01, *p* < 0.001, and *p* < 0.0001 indicated as *, **, ***, and ****, respectively, *ns* indicated no significant differences.

