## supplemental figure legends for "γ-aminobutyrate (GAB) functions as a bioenergetic and signaling gatekeeper to control T cell inflammation"

**Supplemental figure 1. Distinctive extracellular metabolome profiles characterize T cell subsets. a**, Extracellular metabolites in indicated T cell subsets were profiled by LC-MS. Hierarchical clustering heatmap represents the value of the relative amount (see color scale). **b**, Principal component analysis (PCA) for the correlations among each subset. **c**, Pairwise comparison of the statistical analysis, the numbers reflect the correlation R values. **d-f**, Extracellular metabolites associated with energy metabolism (**d**), nucleotide metabolism (**e**), and lipid metabolism (**f**) in indicated T cell subsets were profiled by LC-MS. The value for each metabolite represents the average of triplicates. The complete metabolomic profile is provided in ***Extended information 1***. Statistical analysis was performed by R Programming Language.

**Supplemental figure 2. GAB is an abundant metabolite produced in $T_H$17 and i$T_{reg}$ cells. a**, **b**, GAB in indicated groups was determined and quantified by NMR (*n*=3). **c**, **d**, Indicated metabolites in $T_H$17 cells were quantified by GC-MS. The value for each metabolite represents the average of three samples. The heatmaps (**c**) represent the log2 value (medium) or the absolute value (pellet) of the quantity (see color scale). The complete data is provided in ***Extended information 2***. The volcano plots (**d**) show changes of metabolites in cell pellet (top) and cell medium (bottom). Significance was calculated by unpaired Two-tail Student's *t*-test (**c, d**), data are representative of two independent experiments (**a, b**) or pooled from two experiments (**c, d**). "*n*" indicates the number of biological replicates.

**Supplemental figure 3. Glutamine and Arginine are the main carbon sources for GAB biosynthesis in effector T cells. a**, Schematic diagram of GABA biosynthesis from arginine (top), with the expression of relevant genes determined by qPCR detection (bottom). mRNA levels of $T_{nai}$ were set to 1. The heatmap represents the relative mRNA expression level (see color scale). Values and standard deviations are provided in ***Extended information 3***. **b**, Diagram of [$^{13}C_6$]-glucose conversion to downstream metabolites that are in glycolysis and the TCA cycle. **c**, Indicated metabolites in $T_H$17 cells were quantified by GC-MS (*n*=3). Black dot: $^{12}C$; Red dot: $^{13}C$ derived from [$^{13}C_6$]-glucose. Numbers in the X-axis represent those of $^{13}C$ atoms in given metabolites. Significance was calculated by unpaired Two-tail Student's *t*-test (**c**), data are shown as mean ± SEM, ****$p < 0.0001$. "*n*" indicates the number of biological replicates. Data are representative of three experiments. *ODC*: ornithine decarboxylase, *DAO*: diamine oxidase, PDH: pyrroline dehydrogenase, AG: aminoguanidine, and M: mass spectrum.

**Supplemental figure 4. ABAT enables GAB diverting into the TCA cycle in T cells. a**, **b**, **d**, Diagram of [$^{13}C_4$]-GABA (**a,b,** left) and [$^{13}C_6$]-arginine (**d,** left) conversion to downstream metabolites that are in the TCA cycle and polyamine biosynthesis pathway. Indicated metabolites in $T_H17$ cells were quantified by GC-MS (*n*=3). Black dot: $^{12}C$; Blue dot: $^{13}C$ derived from the indicated tracers. Numbers in the X-axis represent those of $^{13}C$ atoms in given metabolites. **c**, OCR of $T_H17$ cells with indicated treatments was determined by Seahorse. Statistical analysis (*n*=48). Significance was calculated by unpaired Two-tail Student's *t*-test (**a, b, d**), Two-way ANOVA (**c**), data are shown as mean ± SEM, ****$p < 0.0001$. "*n*" indicates the number of biological replicates. Data are representative of three experiments. α-KG: α-Ketoglutarate, OCR: oxygen consumption rate, and M: mass spectrum.

**Supplemental figure 5. ABAT enables GAB diverting into the TCA cycle in T cells. a**, **b**, Diagram of [$^{13}C_4$]-putrescine (**a,** left) and [$^{13}C_5$]-glutamine (**b,** left) conversion to downstream metabolites that are in the TCA cycle and polyamine biosynthesis pathway. Indicated metabolites in $T_H17$ cells were quantified by GC-MS (*n*=3). Black dot: $^{12}C$; Blue dot: $^{13}C$ derived from the indicated tracers. Numbers in the X-axis represent those of $^{13}C$ atoms in given metabolites. **c**, OCR of $T_H17$ cells with indicated treatment was determined by Seahorse. Statistical analysis (*n*=48). Significance was calculated by unpaired Two-tail Student's *t*-test (**a, b**), Two-way ANOVA (**c**), data are shown as mean ± SEM, *$p < 0.05$, **$p < 0.01$, ***$p < 0.001$, ****$p < 0.0001$; *ns,* no significant differences. "*n*" indicates the number of biological replicates. Data are representative of three experiments. *GDH*: glutamate dehydrogenase, OCR: oxygen consumption rate, and M: mass spectrum.

**Supplemental figure 6. ABAT is dispensable for normal T cell development after the double-positive stage. a-f**, Distribution of $CD4^+$ and $CD8^+$ T cell (**a,d**), indicated intracellular proteins (**b,e**) and surface markers (**c,f**) were determined by flow cytometer. Significance was calculated by unpaired Two-tail Student's *t*-test, data are shown as mean ± SEM, *ns,* no significant differences. "*n"* indicates the number of biological replicates.

**Supplemental figure 7. ABAT is dispensable for T cell activation. a**, Cell viability, size and activation markers (**a**), cell cycle profile (**b**), DNA/RNA contents (**c**), protein synthesis activity (**d**), and CFSE dilution and cell viability (**e**) were determined by flow cytometry. *n*=3 in all experiments. Significance was calculated by Two-way ANOVA; data are shown as mean ± SEM, *ns,* no significant differences. "*n"* indicates the number of biological replicates. Data are representative of three experiments. OPP: o-propargyl-puromycin, and MFI: median fluorescence intensity.

**Supplemental figure 8. Inhibition of ABAT suppresses $T_H17$ but enhances $iT_{reg}$ cell differentiation. a-c**, Ingenuity pathway analysis (IPA), gene set enrichment analysis (GSEA), and hierarchical clustering analysis of a list of inflammatory genes were performed by using RNA-seq data of $CD4^+$ T cells that were cultured under $T_H0$ culture condition and collected at 36 hrs after activation (*n*=3). The orange dot-line in X-axis indicates the cut-off value (*p*=0.05) (**a**). The heatmaps (**c**) represent the log2 value of the quantity (see color scale). The complete data is provided in ***Extended information 4***. NES normalized enrichment score, and FDR: false discovery rate.

**Supplemental figure 9. Genetic ablation or pharmacological inhibition of ABAT reduces T cell inflammation in EAE. a-d**, T cells were isolated from indicated sites in experimental animals described in Figure 3e (**a,b**) and Figure 3g (**c,d**). The expression of indicated proteins was determined by flow cytometry. Statistical analysis (*n*=3/ group). Significance was calculated by unpaired Two-tail Student's *t*-test, data are shown as mean ± SEM, $*p < 0.05$, $**p < 0.01$, $***p < 0.001$, $****p < 0.0001$; *ns,* no significant differences. "*n"* indicates the number of biological replicates. Data are pooled from three experiments. Vig: vigabatrin, EAE: experimental autoimmune encephalomyelitis, CNS: central nervous system, LNs: lymph nodes.

**Supplemental Figure 10. Inhibition of ABAT suppresses T cell proliferation and differentiation *in vivo.* a-c,** As illustrated by the experimental diagram of competitive antigen

(OVA)-specific proliferation (**a,** top), the donor cell ratio (**a,** bottom and **b**), CFSE dilution (**a,** bottom), and indicated protein levels (**a,** bottom and **c**), were determined by flow cytometry ($n$=4). **d-e**, As illustrated by the experimental diagram of competitive homeostatic proliferation (**d,** top). The donor cell ratio (**d,** bottom, and **e**) and CFSE dilution (**a,** bottom) were determined by flow cytometry ($n$=3). Significance was calculated by unpaired Two-tail Student's *t*-test, data are shown as mean ± SEM, ** $p < 0.01$, *** $p < 0.001$, ****$p < 0.0001$, *ns,* no significant differences. "*n*" indicates the number of biological replicates. CSA: cervical, submandibular and axilla.

**Supplemental figure 11. GABA receptor is dispensable for normal T cell development after the double-positive stage. a-f**, Distribution of $CD4^+$ and $CD8^+$ T cell (**a,d**), indicated intracellular proteins (**b,e**) and surface markers (**c,f**) were determined by flow cytometer. Significance was calculated by unpaired Two-tail Student's *t*-test, data are shown as mean ± SEM, *ns,* no significant differences. "*n*" indicates the number of biological replicates. Data are representative of two experiments.

**Supplemental Figure 12. GABA receptor is dispensable for regulating T cell activation and proliferation. a-f**,

Cell activation markers (**a,b,e**), CFSE dilution (**c, d, f**) and cell viability (**e,f**) were determined by flow cytometry ($n$=3). Significance was calculated by Two-way ANOVA; data are shown as mean ± SEM, *ns,* no significant differences. "*n*" indicates the number of biological replicates. Data are representative of three experiments. Bicl: bicuculline, PicroT: picrotoxin, Flu: flumazenil.

**Supplemental Figure 13. Genetic ablation of Gabrb3 promotes T cell inflammation in the EAE model. a**,

**a-d**, T cells were isolated from indicated sites in experimental animals described in Figure 4g. The expression of indicated proteins was determined by flow cytometry. Statistical analysis ($n$=3/ group). Significance was calculated by unpaired Two-tail Student's *t*-test, data are shown as mean ± SEM, *$p < 0.05$, **$p < 0.01$, ***$p < 0.001$; *ns,* no significant differences. "*n*" indicates the number of biological replicates. EAE: experimental autoimmune encephalomyelitis, CNS: central nervous system, LNs: lymph nodes.

**Supplemental Figure 14. GAB exerts both bioenergetic and receptor signaling mediated control of T cell differentiation. a,** Schematic conceptual model in which GAB generated by glutamine and arginine can regulate T cell differentiation by entering into the TCA cycle or exporting into the extracellular environment and acting on $GABA_A$ receptor.
