## supplemental figures for "γ-aminobutyrate (GAB) functions as a bioenergetic and signaling gatekeeper to control T cell inflammation"

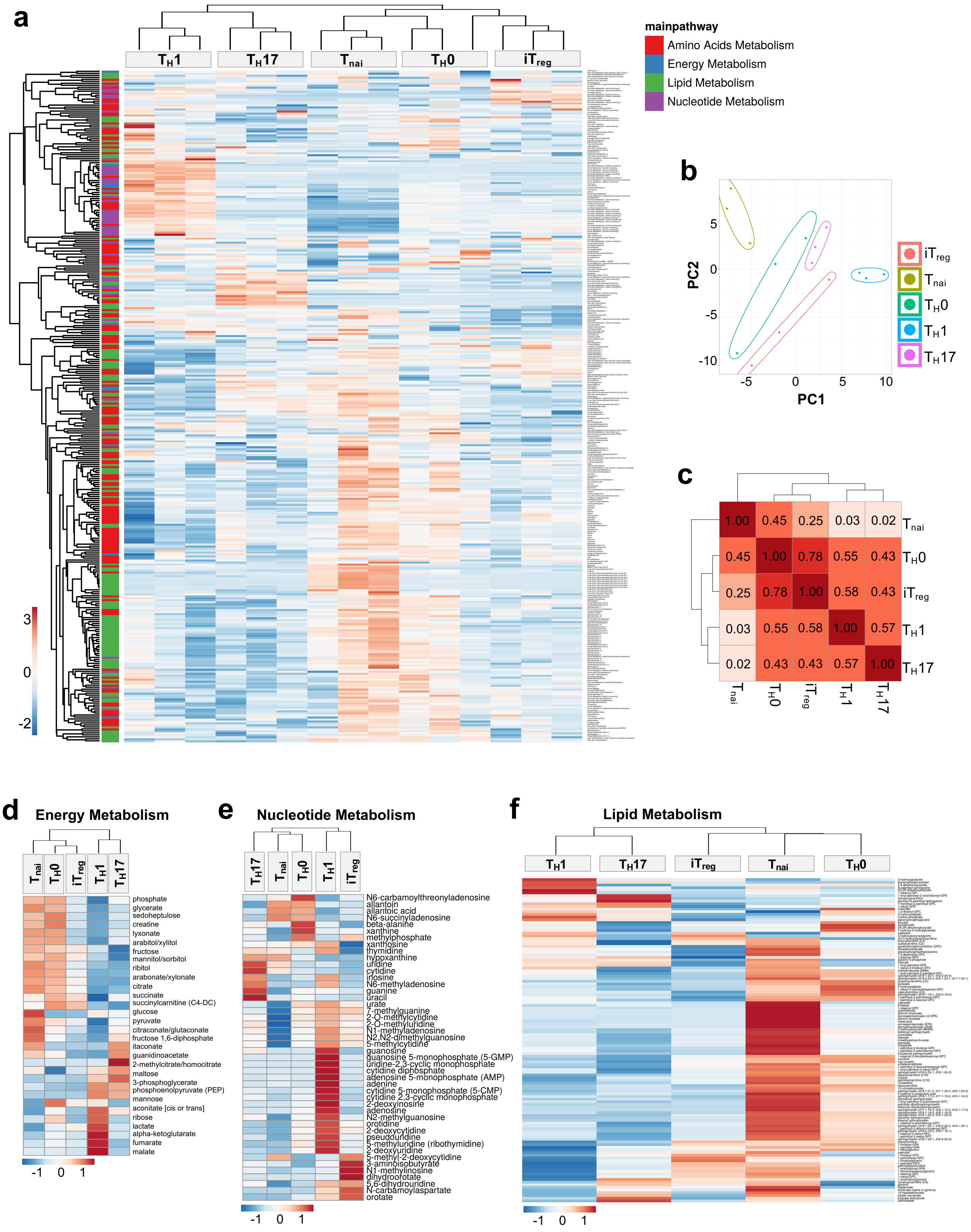


**Supplemental Figure 1. Distinctive extracellular metabolome profiles characterize T cell subsets**

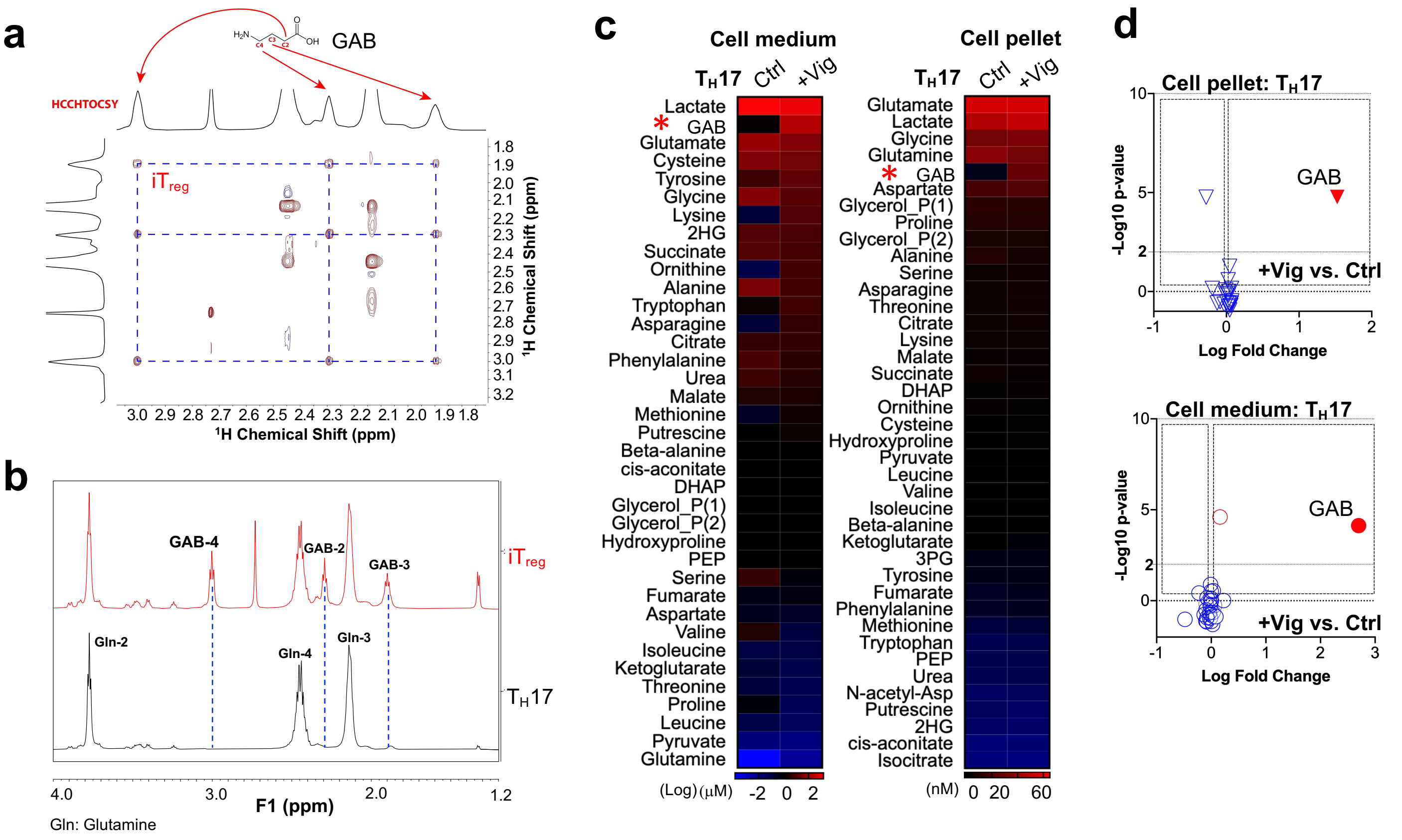


**Supplemental Figure 2. GAB is an abundant metabolite produced in $T_H17$ and $iT_{reg}$ cells**

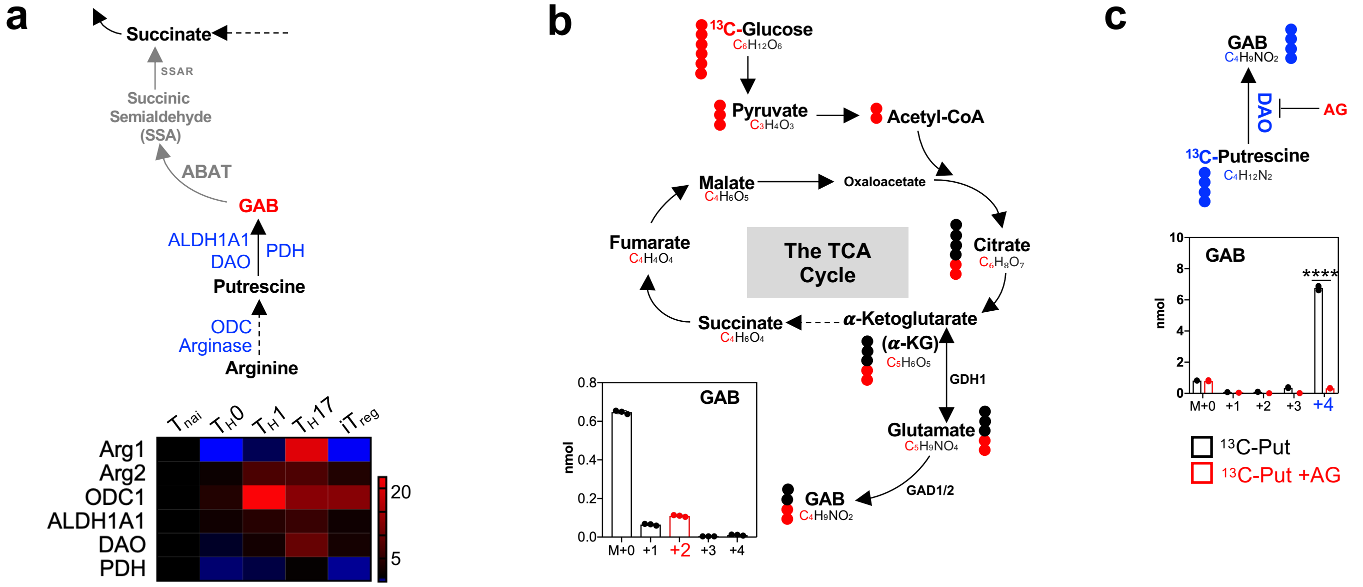


**Supplemental Figure 3. Glutamine and Arginine are the main carbon sources for GAB biosynthesis in effector T cells**

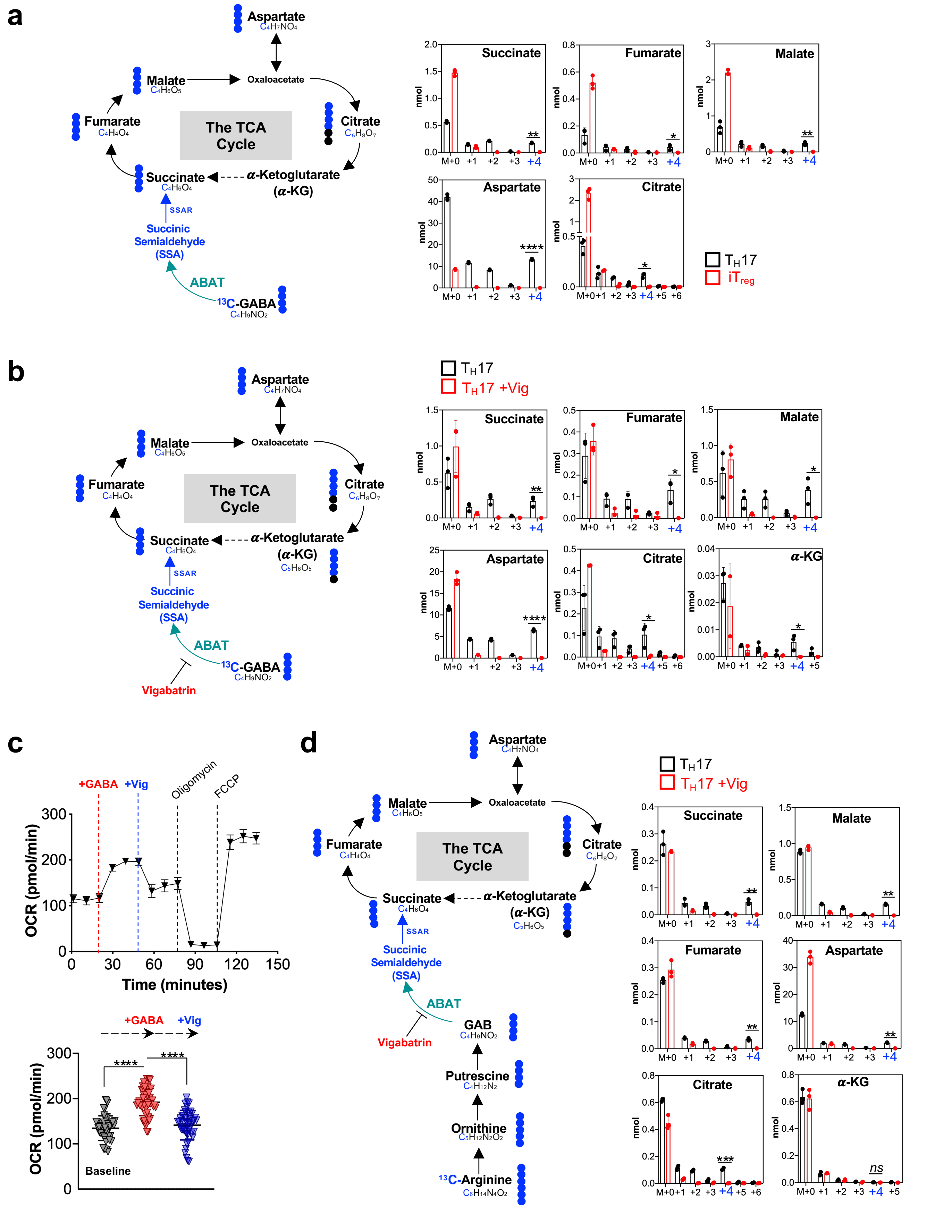


Supplemental Figure 4. ABAT enables GAB diverting into the TCA cycle in T cells

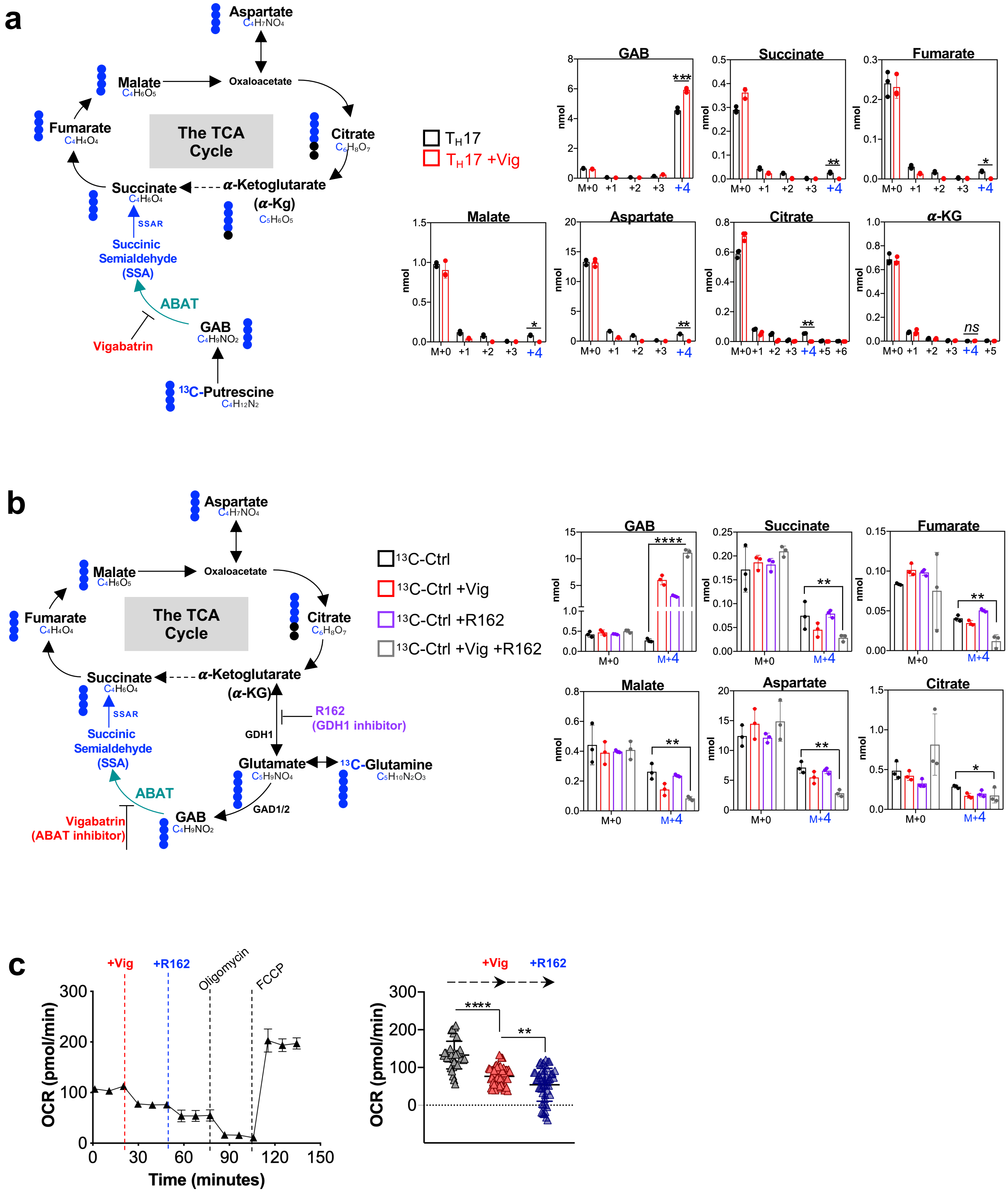


**Supplemental Figure 5. ABAT enables GAB diverting into the TCA cycle in T cells**

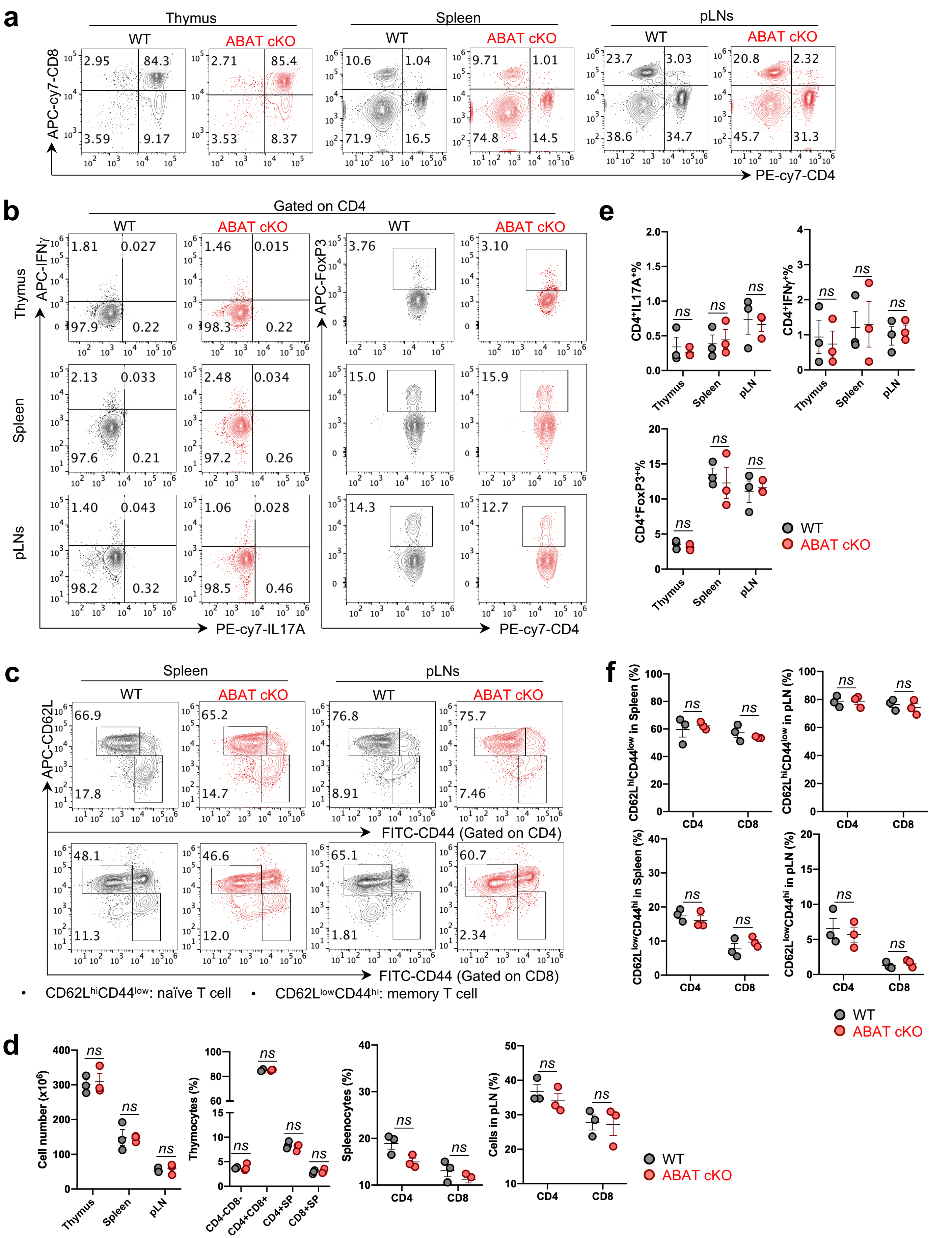


**Supplemental Figure 6. ABAT is dispensable for normal T cell development after the double-positive stage**

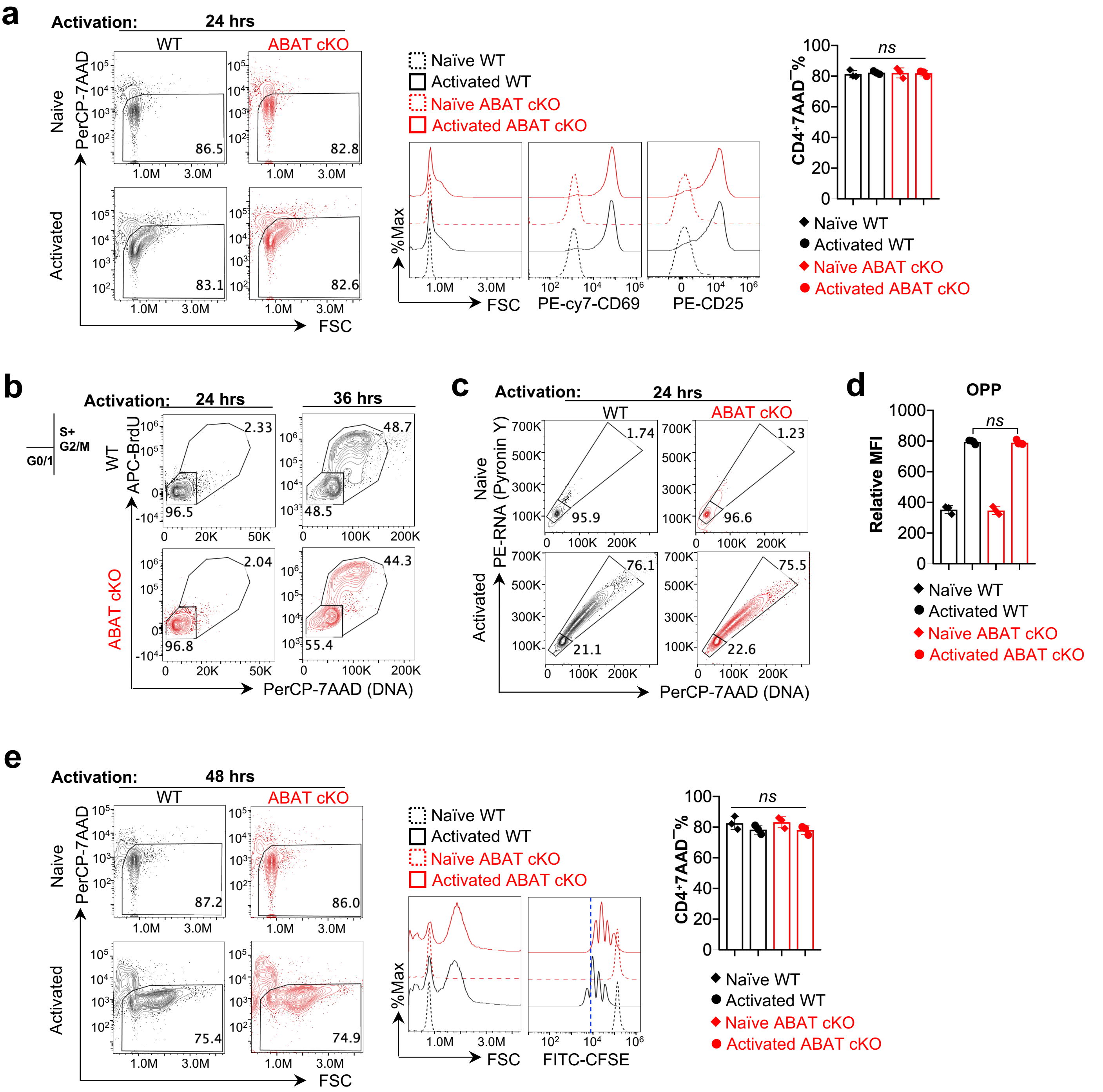


**Supplemental Figure 7. ABAT is dispensable for T cell activation**

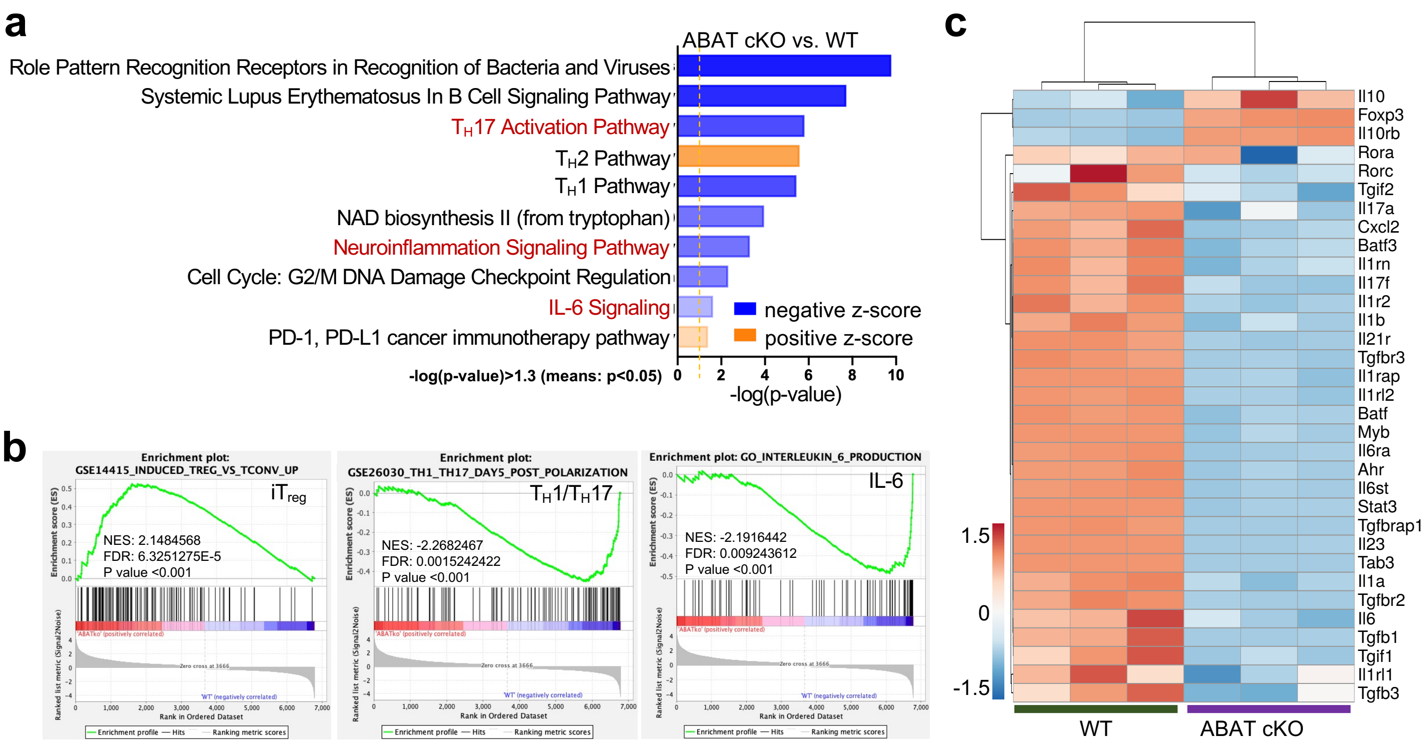


**Supplemental Figure 8. Inhibition of ABAT suppresses $T_H17$ but enhances $iT_{reg}$ cell differentiation**

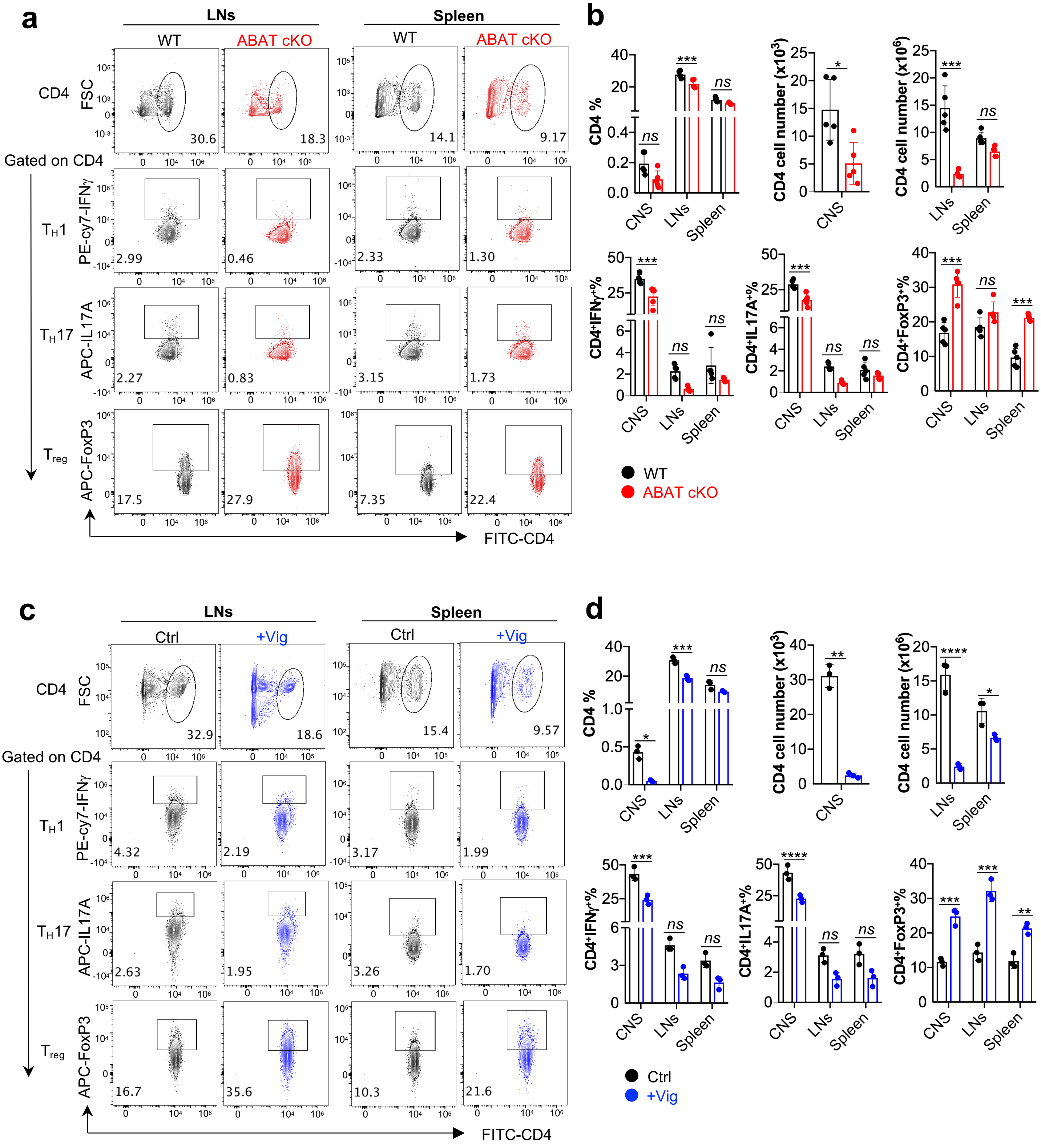

**Supplemental Figure 9. Genetic ablation or pharmacological inhibition of ABAT reduces T cell inflammation in EAE**

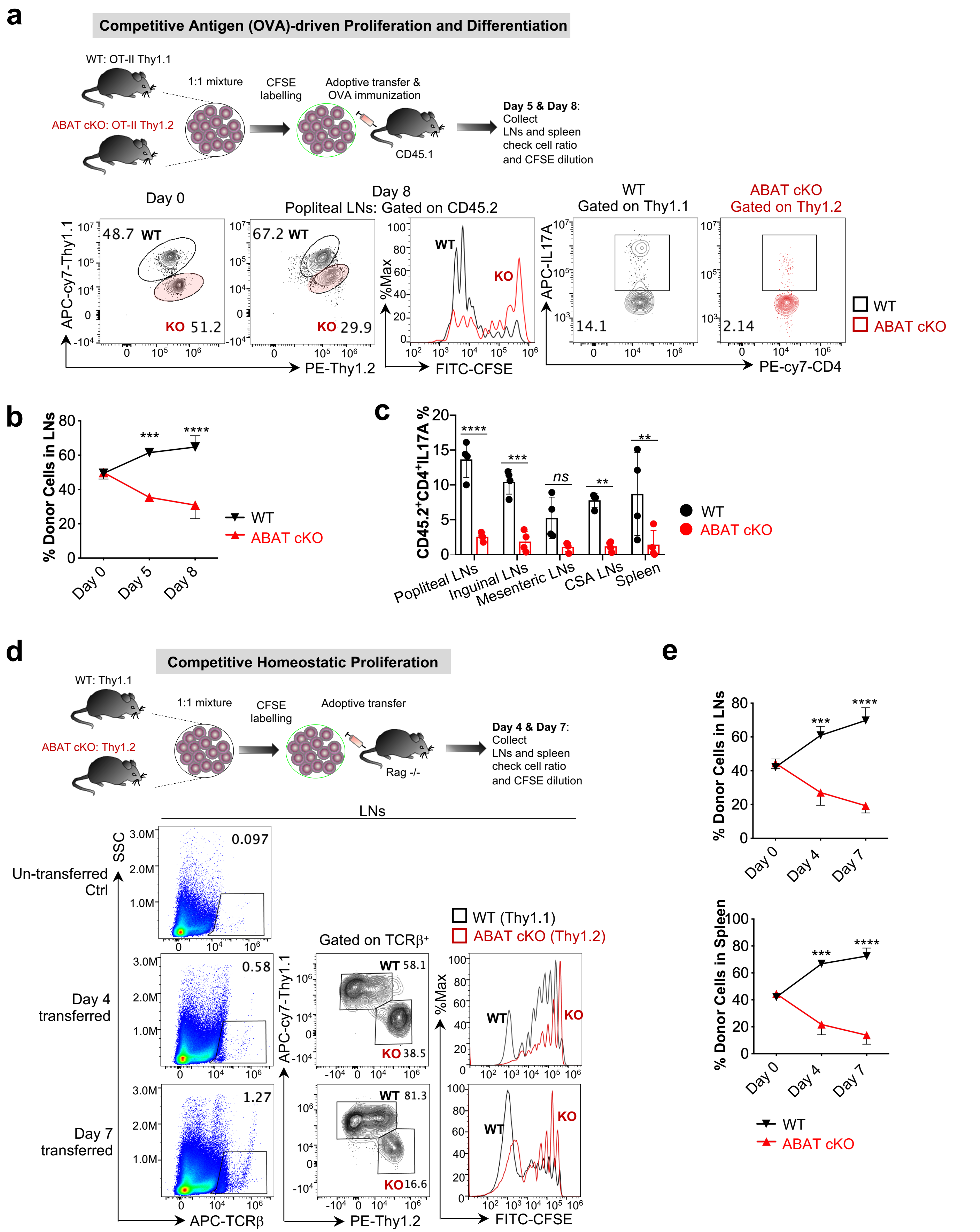


**Supplemental Figure 10. Inhibition of ABAT suppresses T cell proliferation and differentiation *in vivo***

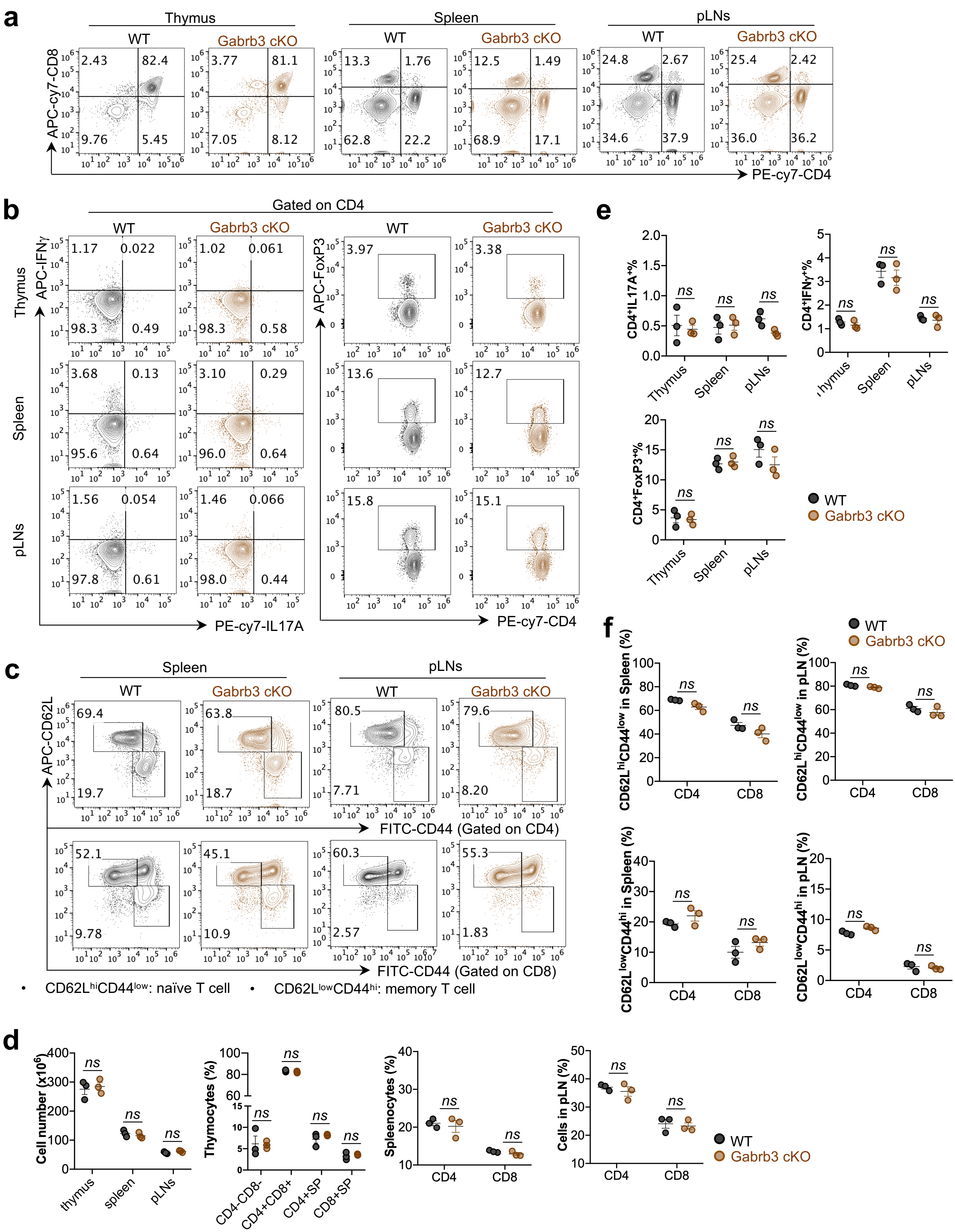


**Supplemental Figure 11. GABA receptor is dispensable for normal T cell development after the double-positive stage**

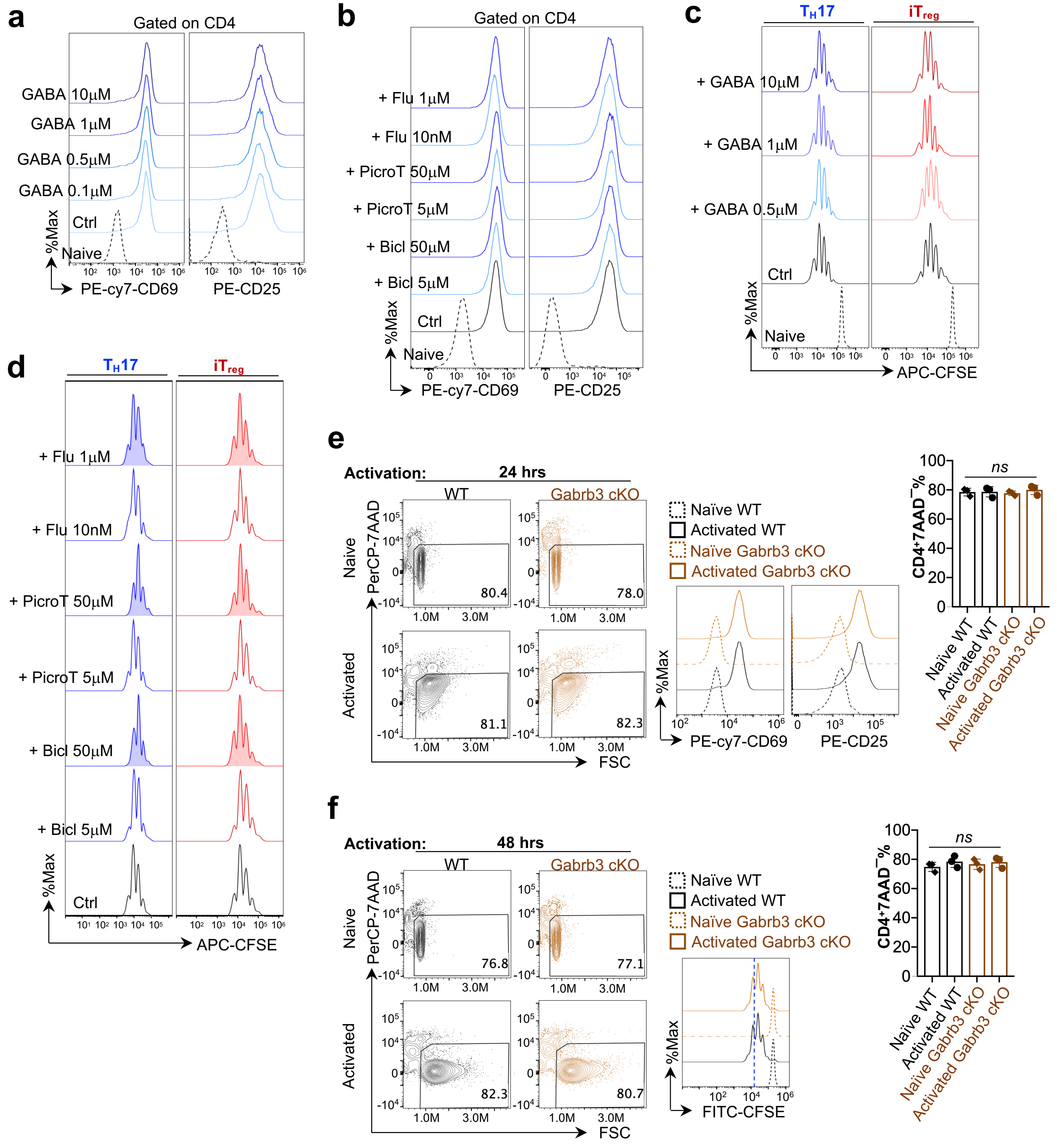


**Supplemental Figure 12. GABA receptor is dispensable for regulating T cell activation and proliferation**

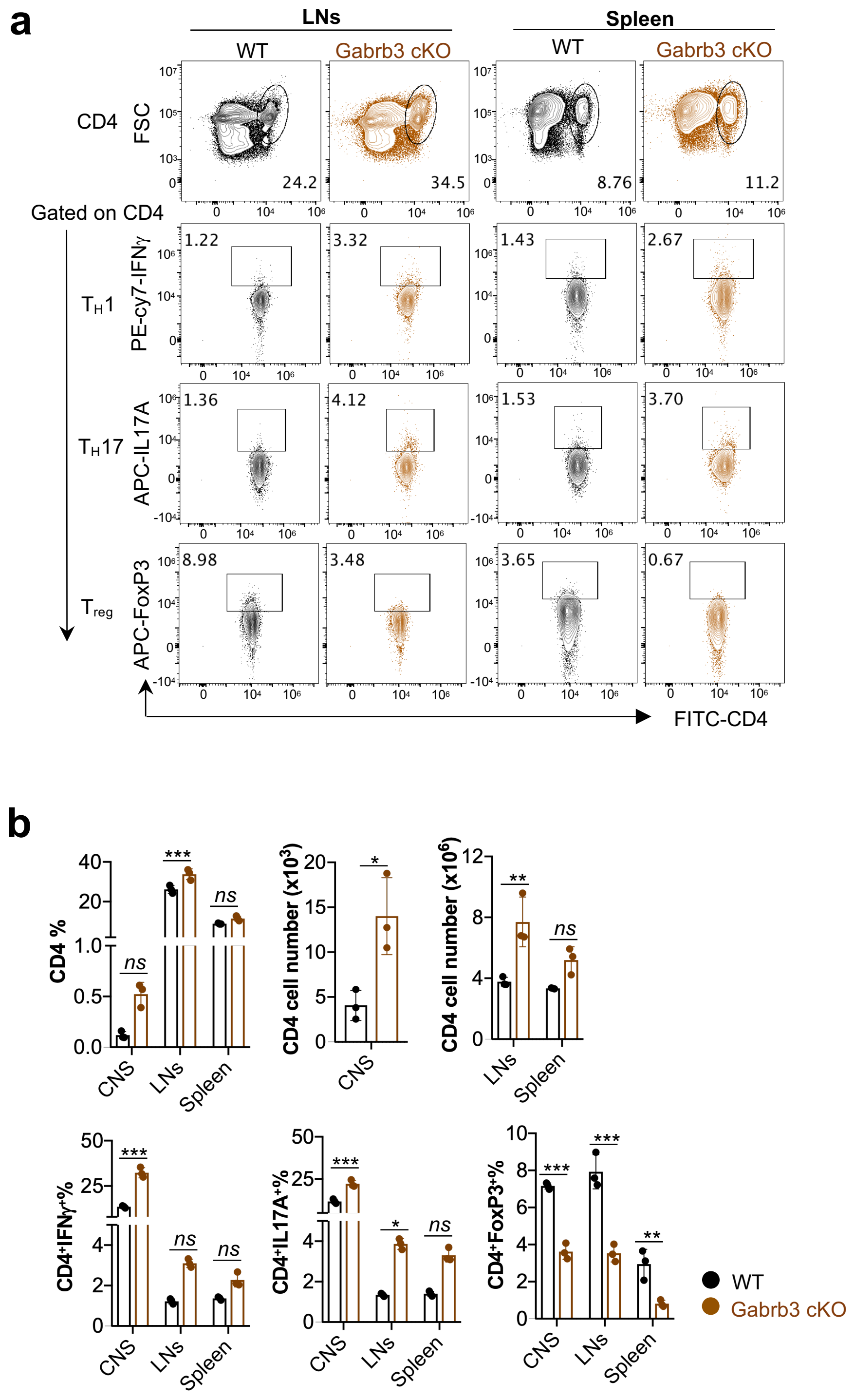


**Supplemental Figure 13. Genetic ablation of Gabrb3 promotes T cell inflammation in the EAE model**

a

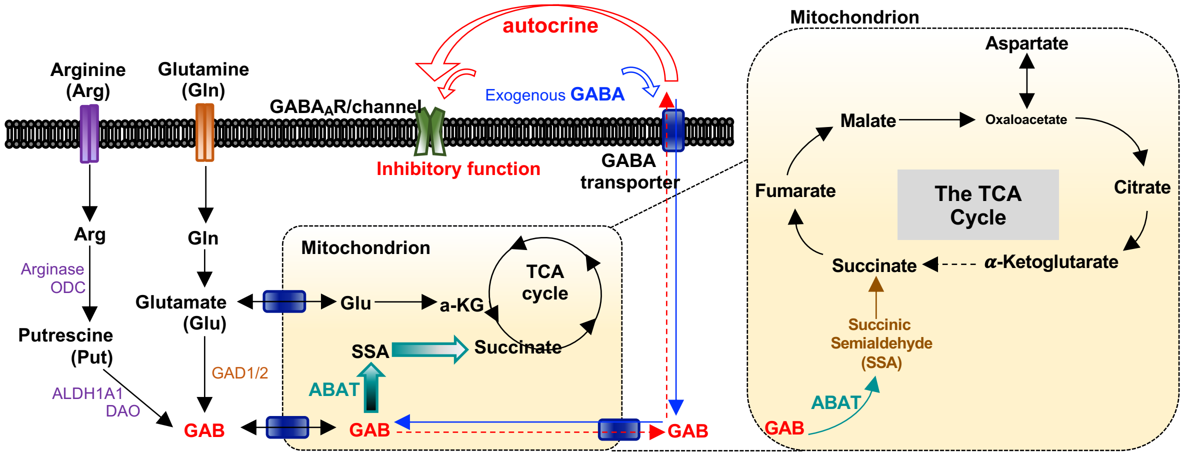


**Supplemental Figure 14. GAB exerts both bioenergetic and receptor signaling mediated control of T cell differentiation**
