## Supplementary material for "γ-aminobutyrate (GAB) functions as a bioenergetic and signaling gatekeeper to control T cell inflammation": tables

**Table 1. Cell culture related antibodies, cytokines, and chemicals**

| Name | Cat# | Vendor | Clone |
|---|---|---|---|
| InVivoMAb anti-mouse CD3 | BE0001-1 | Bio X cell | 145-2C11 |
| InVivoMAb anti-mouse CD28 | BE0015-1 | Bio X cell | 37.51 |
| InVivoMAb anti-mouse IL-2 | BE0043 | Bio X cell | JES6-1A12 |
| InVivoMAb anti-mouse IL-4 | BE0045 | Bio X cell | 11B11 |
| InVivoMAb anti-mouse IFNγ | BP0055 | Bio X cell | XMG1.2 |
| Recombinant Murine IL-12 p70 | 210-12 | PeproTech | |
| Recombinant Human TGF-β1 | 100-21C | PeproTech | |
| Recombinant Murine IL-2 | 212-12 | PeproTech | |
| Recombinant Murine IL-7 | 217-17 | PeproTech | |
| Recombinant Murine IL-6 | 216-16 | PeproTech | |
| g-Aminobutyric acid | A2129 | Sigma | |
| FCCP | C2520 | Sigma | |
| Aminoguanidine; AG | A7009 | Sigma-Aldrich | |
| GDH1 Inhibitor, R162 - Calbiochem | 5.38098 | Sigma-Aldrich | |
| Bicuculline | 11727 | Cayman | |
| Picrotoxin | 20771 | Cayman | |
| Flumazenil | 14252 | Cayman | |
| (R,S)-4-Amino-5-hexenoic acid; Vigabatrin | X-1501 | CNH Technologies | |

**Table 2. Cell staining antibodies and dyes**

| Name | Cat# | Vendor | Clone |
|---|---|---|---|
| FITC anti-mouse CD4 | 100510 | BioLegend | RM4-5 |
| PE/Cyanine7 anti-mouse CD4 | 100422 | BioLegend | GK1.5 |
| APC/Cyanine7 anti-mouse CD8a | 100714 | BioLegend | 53-6.7 |
| PE/Cy7 anti-mouse CD69 | 104512 | BioLegend | H1.2F3 |
| PE anti-mouse CD25 | 102008 | BioLegend | PC61 |
| APC/Cy7 anti-mouse CD90.1 (Thy1.1) | 202520 | BioLegend | OX-7 |
| PE anti-mouse CD90.2 (Thy1.2) | 105308 | BioLegend | 30-H12 |
| APC anti-mouse TCR β chain | 109211 | BioLegend | GL3 |
| PE/Cyanine7 anti-mouse IFN-γ | 505826 | BioLegend | XMG1.2 |
| APC anti-mouse IFN-γ | 505810 | BioLegend | XMG1.2 |
| PE/Cyanine7 anti-mouse IL-17A | 506922 | BioLegend | TC11-18H10.1 |
| APC anti-mouse IL-17A | 506916 | BioLegend | TC11-18H10.1 |
| Alexa Fluor® 647 anti-mouse FOXP3 | 126407 | BioLegend | MF-14 |
| APC anti-mouse CD62L | 104412 | BioLegend | MEL-14 |
| FITC anti-mouse/human CD44 | 103006 | BioLegend | IM7 |
| PerCP anti-mouse CD45.2 | 109826 | BioLegend | 104 |
| FITC anti-mouse/human/rat ABAT | sc-393769 | Santa Cruz Biotechnology | B-12 |
| APC anti-BrdU | 364114 | BioLegend | |
| 7-AAD Viability Staining Solution | 420404 | BioLegend | |
| Pyronin Y | 92-32-0 | Sigma-Aldrich | |

**Table 3. RT-qPCR primers**

| Gene | primer sequences forward | primer sequences reverse |
|---|---|---|
| GAD1 | AACGTATGATACTTGGTGTGGC | CCAGGCTATTGGTCCTTTGTAAG |
| GAD2 | TCCGGCTTTTGGTCCTTCG | ATGCCGCCCGTGAACTTTT |
| ABAT | CTGAACACAATCCAGAATGCAGA | GGTTGTAACCTATGGGCACAG |
| Aldh5a1 | CGGTCAAGGAGAGGAGCTTAC | GGACTAGCCCTCGCTTATCTTT |
| Akr7a5 | CGGCCAGTCCGAGAACATC | TTCAGTGACTTCCCTTCCCAG |
| Glyr1 | GAAACTGGGCCGGTATCCTC | GGTAAGGCTTTAGTTGTTCCACT |
| Slc52a2 | GGGCTCCTACCTGATGACACT | GCACAGGACCCATGAGAGTA |
| Slc6a1 | GAAAGCTGTCTGATTCTGAGGTG | AGCAAACGATGATGGAGTCCC |
| Slc6a11 | TGTTGAGCGTAGCTGGAGAGA | AGCAGATGAAAAACACCACGTA |
| Slc6a12 | GGTCCCTGAGGAAGGAGAGAT | GGGGATGAAGAAAGCTCCACC |
| Slc6a13 | CAGTACACCAACCAGGGAGG | GCCAGGACAACGATGTAGTAGA |
| Slc32a1 | ACCTCCGTGTCCAACAAGTC | CAAAGTCGAGATCGTCGCAGT |
| Gabra1 | TGCCCATGCCTGCCCACTAAAA | GCCATCCCACGCATACCCTCTCT |
| Gabra2 | GGACCCAGTCAGGTTGGTG | TCCTGGTCTAAGCCGATTATCAT |
| Gabra3 | ATGTGGCACTTTTATGTGACCA | CCCCAGGTTCTTGTCGTCTTG |
| Gabra4 | ACAATGAGACTCACCATAAGTGC | GGCCTTTGGTCCAGGTGTAG |
| Gabra5 | TGACCCAAACCCTCCTTGTCT | GTGATGTTGTCATTGGTCTCGT |
| Gabra6 | TGCCCAAGCTCAACTTGAAGA | GCCGTAGACGGTTGTCATAGC |
| Gabarap | AAGAGGAGCATCCGTTCGAGA | GCTTTGGGGGCTTTTTCCAC |
| Gabarapl1 | GGACCACCCCTTCGAGTATC | CCTCTTATCCAGATCAGGGACC |
| Gabarapl2 | TCGGGCTCTCAGATTGTTGAC | ATGGCCTTCTCGGAGGGAA |
| jakmip1 (gababrbp) | ACCGCCTACATCTCGGAACT | GCAGCTCACCTTCCTTGATCTT |
| Gabrb1 | TCCCGTGATGGTTGCTATGG | CCGCAAGCGAATGTCATATCC |
| Gabrb2 | ATGTCGCTGGTTAAAGAGACG | CTGCCACTCGGTTGTCCAAA |
| Gabrb3 | CTGCTGCCAATCTGGCTTTC | CGTAGCCTTTCAACAGCTTGTC |
| Gabrd | ATTGGGGACTACGTGGGCT | CCACATTCACAGGAGCACC |
| Gabre | CCTTCAGTGGAGGGTTGGAC | ATTCAGGCGGAGTTAGAGGCT |
| Gabrg1 | TGTGGAGTCAAACTAGAGGAGTG | TTCCCAGATGCAGGGTTAGTA |
| Gabrg2 | ATGAGTTCGCCAAATACATGGAG | GGAGCAGAATCCACAGCGT |
| Gabrg3 | GACAGTCGCCTTCGATTCAAC | AGCCTCCGCTGTTTTAGAATTT |
| Gabrp | CAGACCCACGGCTAGTGTTC | AGAGGCGGATGAGCCTGTT |
| Gabrq | ATGGGCATCCGAGGTATGCT | ATCCAGAATGACTTCAGGCGT |
| Gabrr1 | CGAGGAGCACACGACGATG | GTGAAGTCCATGTCAACCTCTG |
| Gabrr2 | ATGCCTTATTTGATGAGACTCGC | CCACACCTACAGGGATGGC |
| Gabrr3 | CACCCTAAACGTGAACAACTGT | TCCAATAGTGCCTGAGGTAAAAC |
| Gabbr1 | GCACAGGACACAATGAAAACAG | AGCAAATGTACTCGACTCCCA |
| Gabbr2 | AAGACCCCATAGAGGACATCAA | GGGTGGTACGTGTCTGTGG |
| DBI | GAATTTGACAAAGCCGCTGAG | CCCACAGTAGCTTGTTTGAAGTG |
| Arg1 | CTCCAAGCCAAAGTCCTTAGAG | AGGAGCTGTCATTAGGGACATC |
| Arg2 | CTGCCATTCGAGAAGCTGG | GGGATCATCTTGTGGGACATTAG |
| ODC1 | GGTTCCAGAGGCCAAACA | CAGCGTGCCATCATCCT |
| Aldh1a1 | ATACTTGTCGGATTTAGGAGGCT | GGGCCTATCTTCCAAATGAACA |
| DAO | GGTGGCAAGAGGAGTGGATG | TGGGATGATGTACGGAGAGTTG |
| Aldh4a1 (PDH) | CGATGGAAGCACACCTCTTCT | GGCGACAACTGGTACTGTATATC |
| Tubulin | TTCTGGTGCTTGTCTCACTGA | CAGTATGTTCGGCTTCCCATTC |

**Table 4. Stable isotope tracers**

| Name | Cat# | Vendor |
|---|---|---|
| U13C5-Arginine | CLM2265 | Cambridge Isotope Laboratories, Inc. |
| U13C6-Glucose | CLM1396 | |
| U13C5-Glutamine | CLM1822 | |
| 4-Aminobutyric acid (GABA)(13C4, 97-99%) | CLM-8666 | |
| 1,4-BUTANEDIAMINE (PUTRECINE) (13C4, 98%) | CLM-6574 | |
